# NLRC5 expression in tumor cells is critical to activate adaptive and innate antitumor immune responses

**DOI:** 10.64898/2025.12.19.695247

**Authors:** Akhil Shukla, Akouavi Julite Irmine Quenum, Jean-François Lucier, Anny Armas Cayarga, Dominique Lévesque, François-Michel Boisvert, Thomas A. Kufer, Sheela Ramanathan, Subburaj Ilangumaran

## Abstract

Tumors evade cytotoxic T lymphocyte (CTL)-mediated killing by downregulating MHC class-I, mainly resulting from the loss of its transcriptional activator NLRC5. Expressing full-length NLRC5 (NLRC5-FL) or a shorter NLRC5-CIITA fusion protein termed NLRC5 super-activator (NLRC5-SA) in cancer cells upregulates MHC-I expression and promotes antitumor immunity. To distinguish the role of NLRC5 expressed within tumor cells and antigen presenting cells, we studied B16-F10 melanoma expressing NLRC5-FL (B16-N-FL) or NLRC5-SA (B16-N-SA) in *Nlrc5^+/+^* and *Nlrc5^−/−^* mice. Both tumors were efficiently controlled in both *Nlrc5^+/+^* and *Nlrc5^−/−^* hosts with abundant immune cell infiltration, enriched for activated and differentiated CD8^+^ and CD4^+^ T cells, NK, NKT and iNKT cells. B16-N-FL and B16-N-SA tumors showed increased collagen deposition and vascularization, with upregulation of CCL4 and CXCL9 chemokine genes in B16-N-SA tumors. Depletion of either CD8^+^ T cells or NK1.1^+^ cells increased the growth of B16-N-FL and B16-N-SA tumors in *Nlrc5^+/+^* mice, and that of B16-N-SA tumors in *Nlrc5^-/-^* hosts. Proteomes of B16-N-FL and B16-N-SA cells showed downmodulation of dominant tumor antigens and upregulation of ubiquitination and protein processing pathway proteins. Differentially expressed proteins shared between B16-N-FL and B16-N-SA cells showed enrichment in phagosome and autophagy pathways. We conclude that tumor cell-intrinsic NLRC5 expression is critical for the activation of adaptive and innate immune cells, and establishment of an immune-supportive tumor microenvironment to permit immune cell infiltration and their effector functions and achieve tumor control. NLRC5 expression in APCs is dispensable to mediate these effects. Delivering NLRC5-SA is a promising approach to restore antitumor immune responses in MHC-I-low immune evasive tumors.

**Graphical abstract:** 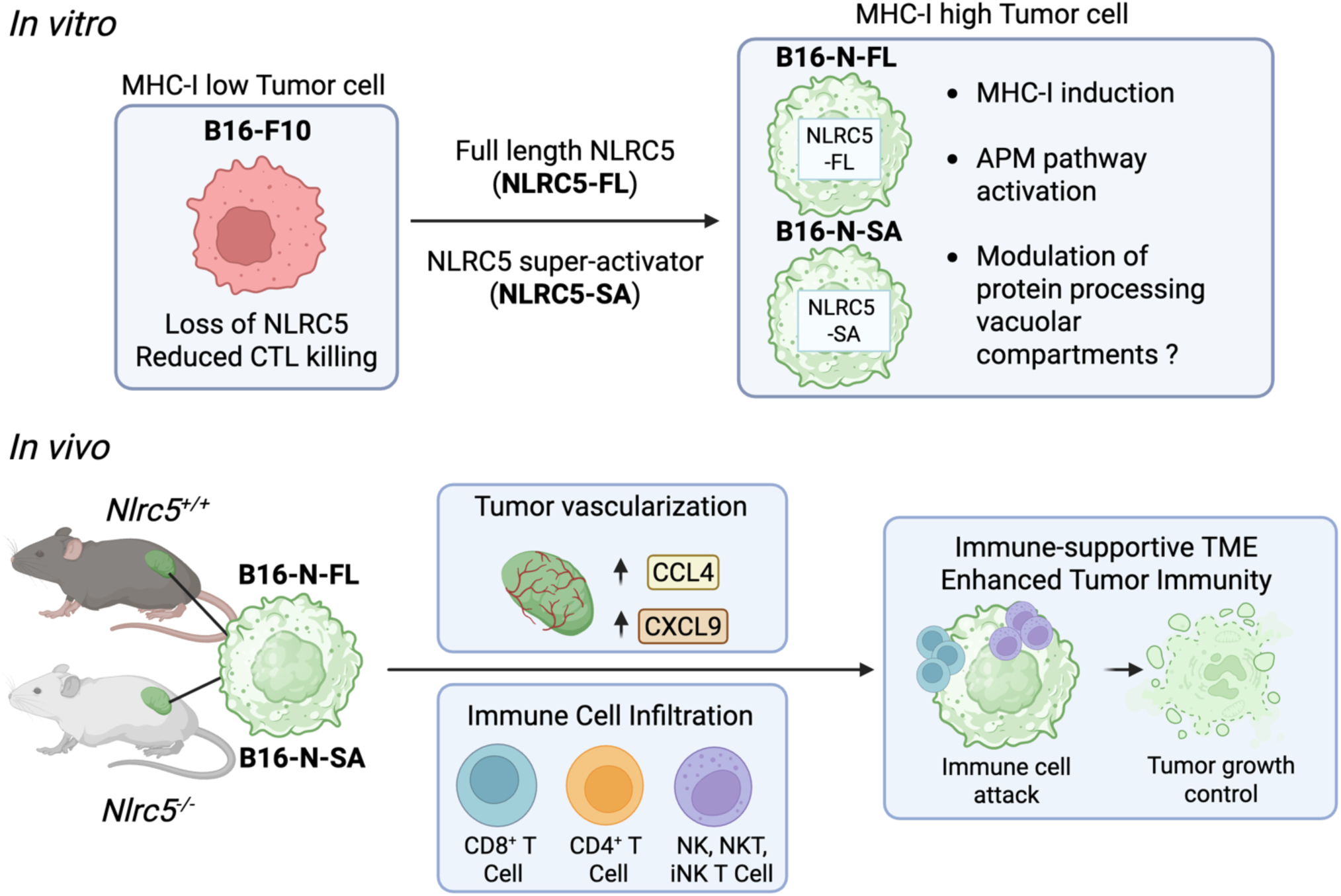

## Introduction

A central cancer immune evasion mechanism is the downmodulation of major histocompatibility class-I (MHC-I) and antigen processing machinery (APM) genes ^1,2^. This defect impairs tumor antigen presentation, renders tumors less immunogenic and allows them to evade killing by CD8^+^ cytotoxic T lymphocytes (CTL). Loss of MHC-I also impairs responsiveness to immune checkpoint blockade (ICB) therapy ^3,4^. Restoring MHC-I expression in cancers is considered a promising way to increase tumor immunogenicity and achieve tumor control but this goal remains challenging ^5–7^.

Downmodulation of MHC-I in tumors results mainly from the loss of NLRC5 (*NOD-Like Receptor CARD domain containing 5*) expression ^8^. NLRC5 is the master transcriptional activator for MHC class-I genes and several antigen processing machinery (APM) genes ^9^. We have previously shown that expressing NLRC5 in MHC-I-low tumors upregulates MHC-I and APM genes, promotes tumor immunogenicity and elicits CD8^+^ T cell-dependent protective antitumor immunity ^10^. Reduced NLRC5 expression in tumors correlates with diminished CTL infiltration and unresponsiveness to ICB therapy ^11,12^. Using 3’-methylcholanthrene-induced fibrosarcoma model in NLRC5 deficient mice, we have recently reported that NLRC5 is required for efficient cancer immune surveillance and cancer immunoediting by lymphocytes ^13^.

The above studies raised the possibility of exploiting NLRC5 for cancer immunotherapy against MHC-I-low immune evasive cancers ^14–16^. As the large molecular size of full-length NLRC5 (NLRC5-FL) restrains its translational utility, we tested a smaller engineered NLRC5-CIITA construct that is as efficient as NLRC5-FL in inducing MHC-I expression and thus designated as NLRC5 super-activator (NLRC5-SA) ^15,17^. MHC-I-associated peptides eluted from EL4 thymoma cells expressing NLRC5-SA or NLRC5-FL showed overlapping and distinct peptides, suggesting differences in tumor antigen processing and presentation by NLRC5-FL and NLRC5-SA ^15^. Nonetheless, NLRC5-SA was as efficient as NLRC5-FL in controlling EL4 tumors and B16-F10 melanoma, supporting its potential application in immunotherapy against MHC-I-low immune evasive cancers ^15^.

In the cancer immunity cycle, activation of naïve CD8^+^ T cells reactive to tumor antigens occurs in tumor draining lymph nodes by antigen presenting cells (APC) ^18^. Activated CD8^+^ T cells proliferate and differentiate into effector CTLs, enter the circulation, traffic to the tumor, recognize antigenic peptides on cancer cells and release cytotoxic granules to eliminate them ^18^. The release of more tumor antigens and their presentation to naïve CD8^+^ T cells, and iteration of this cycle results in tumor control. Thus, loss of NLRC5 expression and downmodulation of MHC-I in cancers will not only impair tumor cell killing but also compromise the tumor immunity cycle and restrain the breadth of anti-tumor CTL response. Dendritic cells (DC) acquire tumor antigens from dead tumor cells and cross-present tumor antigenic peptides on their own MHC-I molecules along with costimulatory ligands ^19,20^. APCs can also acquire MHC-I-peptide complexes directly from tumor cells via trogocytosis and present them to CD8^+^ T cells by cross-dressing ^21,22^. The impact of the loss of NLRC5 in cancer cells on presentation of tumor antigens by APCs has not been yet addressed.

In addition to T cells, innate immune cells such as NK cells also contribute to tumor immune surveillance ^23,24^. NK and NKT cells kill cancer cells by recognizing the absence of self-MHC-I molecules. In addition to upregulating classical MHC-Ia molecules, NLRC5 induces non-classical MHC-Ib molecules ^25–28^, which are recognized by activating and inhibitory receptors expressed on NK cells ^29–31^. As NLRC5 deficiency compromises the expression of both classical and non-classical MHC-I molecules, it is unclear why tumors that downregulate NLRC5 are not controlled by innate immune cells. To investigate how NLRC5 expression in tumor cells contributes to the activation of adaptive and innate cells, and to determine the role of endogenous NLRC5 in host APCs in antitumor immune responses, here we studied tumor formation by B16-F10 melanoma cells expressing NLRC5-FL or NLRC5-SA in NLRC5-deficient and control mice and evaluated the associated cellular and molecular mechanisms. Our findings show that NLRC5 expression within tumor cells is necessary and sufficient for efficient activation of adaptive and innate immune cells and for achieving effective control of MHC-I- low immune evasive cancers.

## Results

### NLRC5 expression in tumor cells is required for effective immune-mediated tumor control

B16-F10 melanoma cells stably expressing NLRC5 (B16-N-FL), an engineered version designated as NLRC5 super-activator (B16-N-SA) or the control vector (B16-V) were first assessed for growth kinetics in vitro. All three cell lines showed comparable levels of viability and proliferation at 24, 48 and 72 hours, indicating that stable expression of NLRC5-FL or NLRC5-SA did not affect the growth of B16 cells (Supplementary Fig. S1A). This was confirmed by the comparable growth of B16-V, B16-N-FL and B16-N-SA cell lines as subcutaneous tumors in immunodeficient NSG mice, wherein all tumors reached the endpoint within 14 days (Supplementary Fig. S1B, C). Upon implantation into immunocompetent C57Bl/6 mice, tumors formed by B16-V cells reached only half the size of B16-V tumors growing in NSG mice at 14 days post-implantation, indicating that a functional immune system restrains the growth of B16-V cells (Supplementary Fig. S1D) despite their low MHC-I expression ^32^. B16-N-FL and B16-N-SA cells, which show elevated expression of MHC-I ^10,15^, formed significantly smaller tumors than B16-V tumors in C57Bl/6 mice (Supplementary Fig. S1C, D). These data support the potential utility of NLRC5-SA in restoring antitumor immunity against MHC-I-low tumors, and that a greater understanding of the antitumor functions of NLRC5-SA would strengthen its exploitation for cancer immunotherapy.

Adaptive antitumor immune response is initiated following the priming of naïve antitumor CD8^+^ T cells by professional antigen presenting cells (APCs) in tumor draining lymph nodes through cross presentation of tumor antigens ^18^. To distinguish the impact of NLRC5 expressed in tumor cells from that of endogenous NLRC5 expressed in APCs on antitumor immunity, we compared the growth of B16-N-FL and B16-N-SA tumors in *Nlrc5^−/−^* and littermate *Nlrc5^+/+^* hosts. Both tumors were efficiently controlled in *Nlrc5^+/+^* hosts (Fig. 1A,C-D), as expected ^15^. Notably, *Nlrc5^−/−^*mice were as efficient as *Nlrc5^+/+^* hosts in controlling B16-N-FL and B16-N-SA tumors (Fig. 1B-D). B16-V tumors also showed discernibly slower growth in *Nlrc5^−/−^* hosts. Histological examination revealed that tumors formed by B16-N-FL and B16-N-SA cells contained prominent areas of necrosis compared to B16-V tumors in both *Nlrc5^+/+^* and *Nlrc5^−/−^* hosts (Fig. 1E, F). These necrotic areas are unlikely to arise from cell death caused by nutrient and oxygen deprivation resulting from tumor growth because such lesions were not evident in much larger B16-V tumors. Alternatively, these necrotic lesions could be caused by immune cell-mediated killing of tumor cells. This possibility is supported by the significantly elevated infiltration of CD45^+^ immune cells in B16-N-FL and B16-N-SA tumors in *Nlrc5^+/+^* mice and in B16-N-SA tumors of *Nlrc5^−/−^*hosts compared to B16-V tumors (Fig. 1G, H). B16-V tumors in *Nlrc5^−/−^*hosts contained discernibly more CD45^+^ cells, although this difference was not statistically significant. These results indicate that NLRC5 expression within tumor cells is necessary and sufficient, whereas endogenous NLRC5 expressed in host APC is dispensable, to promote immune cell infiltration and achieve efficient tumor control, which can be achieved by delivering NLRC5-SA to tumor cells.

**Figure 1.**
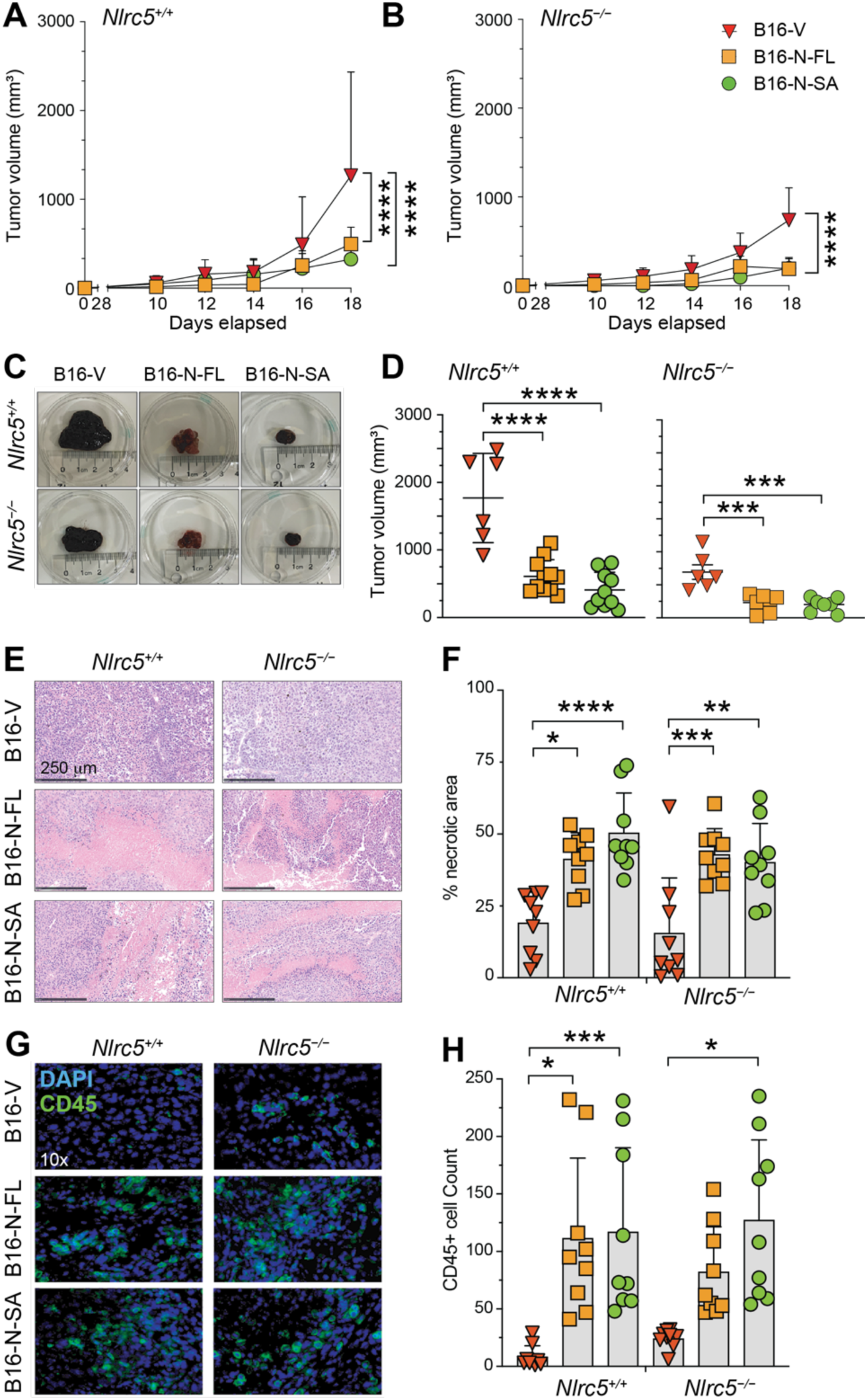
NLRC5 expression within tumor cells is necessary and sufficient for tumor control by immune cells. B16-F10 melanoma cells (2 ×10^5^ cells in 50 μL PBS) expressing control vector (B16-V), full length NLRC5 (B16-N-FL) or engineered NLRC5 super-activator (B16-N-SA) were implanted via subcutaneous route in the right flank of *Nlrc5^+/+^* and *Nlrc5^−/−^*littermate controls. Tumor growth was measured every 2 days from day 8 onwards. When B16-V tumor reached the endpoint (20 mm in diameter in any one direction) in more than one of the recipient mice, all mice were euthanized, and the tumor tissues were processed for paraffin embedding. (A, B) Tumor growth kinetics in littermate *Nlrc5^+/+^* and *Nlrc5^−/−^*hosts. Tumor volume was (volume calculated using ellipsoid formula [½*length*(width^2^)]). Data shown from 6-10 mice per group from two independent experiments. (C) Representative images of tumors dissected from *Nlrc5^+/+^* and *Nlrc5^−/−^* hosts at the endpoint (using consistent zoom factor of 1.7x). (D) Tumor mass at the endpoint in *Nlrc5^+/+^* and *Nlrc5^−/−^* hosts. (E) Hematoxylin and eosin stained tissues of representative tumors. Scale bar, 250 μm. * indicates areas of necrosis. (F) Quantification of the necrotic areas. The proportion of necrotic area was calculated from nine random fields from three tumors per group using ImageJ. (G) Immunofluorescence staining of CD45 marker in representative tumors. Magnification 10×. (H) Quantification of CD45^+^ cells from nine random fields areas with a minimum of 100 cells per field. (A,B,D,F,H) Statistics: Mean + Standard deviation (SD). Ordinary one-way ANOVA with Tukey’s multiple comparison test. * *p* ≤0.05, ** *p* ≤0.01, *** *p* ≤0.001, **** *p* ≤0.0001.

### Tumor-intrinsic NLRC5 expression promotes T cell and NK cell infiltration

To characterize the immune cells associated with the control of B16-N-FL and B16-N-SA tumors, we characterized tumor infiltrating lymphocytes (TILs) in B16-V, B16-N-FL and B16-N-SA tumors isolated from *Nlrc5^+/+^* and *Nlrc5^−/−^* hosts at the endpoint. In parallel, we studied the immune cell profiles in tumor draining (DLN) and non-tumor draining (NDLN) lymph nodes collected simultaneously from these mice (Fig. 2). As the tumors were different in size, the number of CD45^+^ cells and lymphocyte subsets were normalized for the tumor mass. The numbers of CD45^+^ cells per gram of tissue were significantly elevated in B16-N-FL tumors compared to B16-V tumors in *Nlrc5^+/+^* and *Nlrc5^−/−^*mice, whereas B16-N-SA tumors harbored more CD45^+^ cells only in *Nlrc5^−/−^* mice (Fig. 2A). On the other hand, B16-N-SA tumors caused an increase in the cellularity of DLN in both *Nlrc5^+/+^* and *Nlrc5^−/−^*hosts, whereas B16-N-FL increased cell numbers in the DLN of *Nlrc5^−/−^* mice (Fig. 2A). The cellularity of NDLN was not altered by B16-N-FL or B16-N-SA tumors in either host.

**Figure 2.**
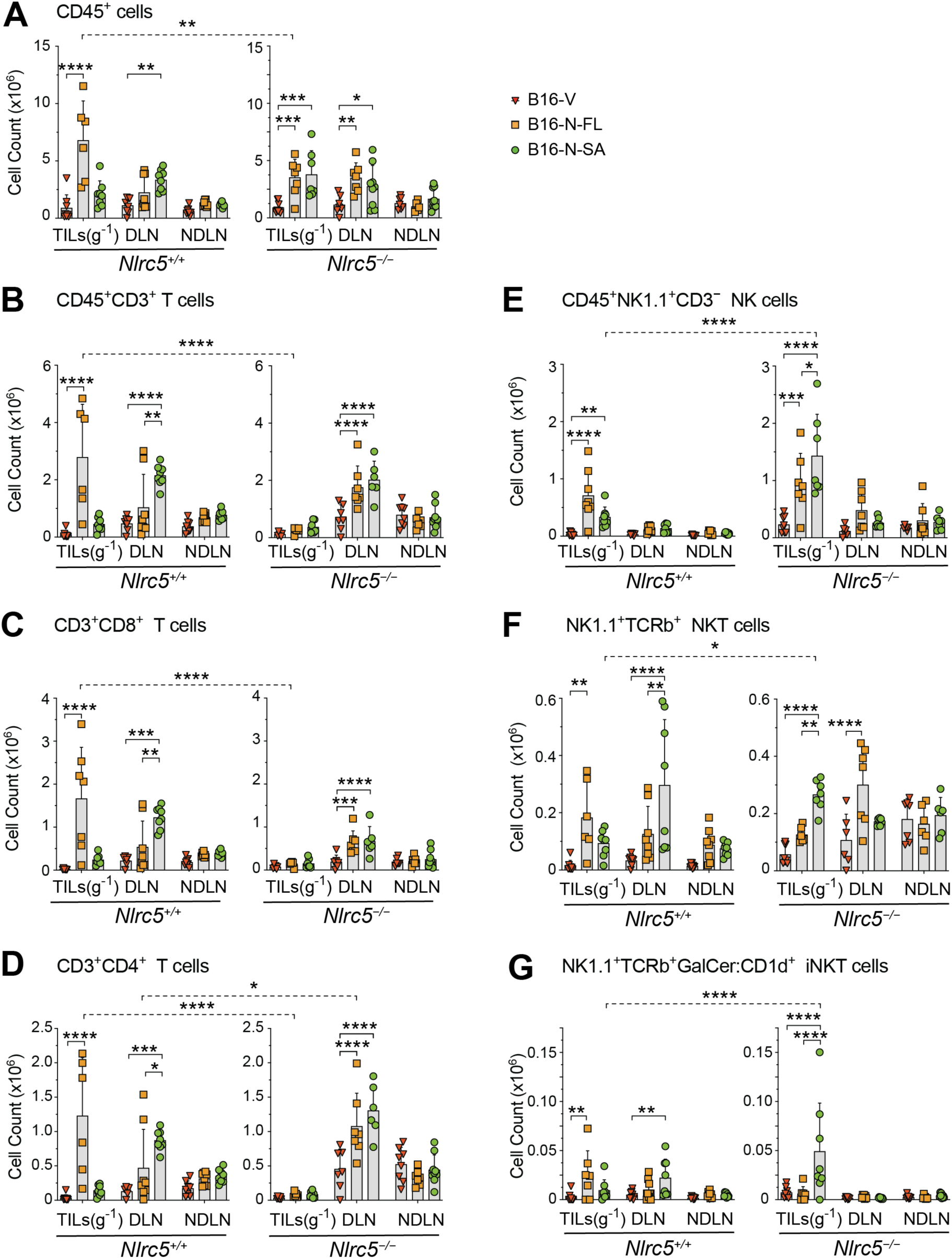
NLRC5-FL or NLRC5-SA expression in tumor promotes lymphoid and NK cell infiltration in *Nlrc5^+/+^*and *Nlrc5^−/−^* hosts. Tumor infiltrating lymphocytes (TILs) and single cell suspensions from tumor draining (DLN) and non-tumor draining (NDLN) lymph nodes were isolated at the time of tumor collection from *Nlrc5^+/+^* and *Nlrc5^−/−^* hosts implanted with B16-V, B16-N-FL or B16-N-SA cells. Cells were counted and stained with fluorochrome conjugated antibodies for lymphoid and NK cell markers and analyzed by flow cytometry using gating strategies depicted in Supplementary Fig. S2. Data from 6-9 mice per group from two independent experiments are shown. (A) Total cellularity (number of CD45^+^ singlet cells) in TILs, DLN and NDLN. The cell numbers in TILs were equalized to the weight of the tumor mass (TILs per gram; TILS(g^-1^)). (B-G) Absolute number of total CD3^+^ T cells (B), CD4^+^ T cells (C), CD8^+^ T cells (D), NK cells (E, NK1.1^+^TCRb^−^), NKT cells (F, NK1.1^+^TCRb^+^) and iNKT cells (G, NK1.1^+^TCRb^+^ α-GalCer:CD1d^+^) were calculated from their percentages within the gated populations (Supplementary Fig. S3). (A-G) Statistics: Mean + SD. Two-way ANOVA with Tukey’s multiple comparison test. * *p* ≤0.05, ** *p* ≤0.01, *** *p* ≤0.001, **** *p* ≤0.0001. Significant differences between B16-V, B16-N-FL and B16-N-SA are indicated by solid lines. Significant differences between *Nlrc5^+/+^*and *Nlrc5^−/−^* hosts for the same tumor are indicated by dotted lines.

Immunophenotyping of TILs, DLN and NDLN cells was performed following the gating strategies shown in Supplementary Fig. S2. The absolute numbers of various lymphoid cell subsets were calculated from their proportions, shown in Supplementary Fig. S3. TILs of B16-N-FL tumors harbored significantly more CD3^+^ T cells and CD4^+^ and CD8^+^ T cell subsets than B16-V tumors in *Nlrc5^+/+^* but not in *Nlrc5^−/−^* hosts, whereas these cell numbers were not increased in the TILs of B16-N-SA tumors in either of the hosts (Fig. 2B, C, D). On the other hand, total CD3^+^ and CD8^+^ and CD4^+^ T cell numbers were significantly elevated in the DLN of B16-N-SA tumors compared to B16-V tumors in both *Nlrc5^+/+^* and *Nlrc5^−/−^* hosts, whereas an increase in these cell numbers in the DLN of B16-N-FL tumors occurred only in *Nlrc5^−/−^* mice (Fig. 2B, C, D). NK cell subsets displayed differential enrichment in the TILs and DLN of B16-N-FL and B16-N-SA tumor bearing mice. NK1.1^+^CD3^−^ NK cells were significantly enriched in the TILs of B16-N-FL and B16-N-SA tumors in both *Nlrc5^+/+^* and *Nlrc5^−/−^*hosts, but their numbers did not increase in DLN (Fig. 2E). NK1.1^+^ TCRb^+^ NKT cells and NK1.1^+^α-GalCer:CD1d^+^ iNKT cells were also significantly enriched in TILs of B16-N-FL tumors in *Nlrc5^+/+^* mice, whereas B16-N-SA tumors showed their enrichment only in TILs from *Nlrc5^−/−^* hosts and in the DLN of *Nlrc5^+/+^* hosts (Fig. 2F, G).

These data indicate that NLRC5-FL and NLRC5-SA expressing tumors promote the activation of T cells in DLN and the recruitment of T, NK, NKT and iNKT cells into tumors, independent of NLRC5 expression in host APCs. While NLRC5-SA appears to be more effective than NLRC5-FL in recruiting T, NKT and iNKT cells in DLN, the enrichment of NK cell subsets in TILs of *Nlrc5^−/−^* hosts compared to *Nlrc5^+/+^* hosts suggests that endogenous NLRC5 expressed in host cells likely modulates NK, NKT and iNKT cell activation and antitumor functions.

### NLRC5 expression in tumor cells promotes intratumoral T cell activation

As NLRC5 upregulates MHC-I and tumor antigen presentation ^10^, the activation and differentiation of CD8^+^ T cells and their infiltration into tumors expressing NLRC5-FL or NLRC5-SA were examined. TILs from B16-N-FL and B16-N-SA tumors and their DLN harbored significantly elevated numbers of CD69^+^ activated effector (T_E_) CD8^+^ T cells compared to B16-V tumors in both *Nlrc5^+/+^* and *Nlrc5^−/−^* hosts (Fig. 3A). B16-N-FL and B16-N-SA TILs were similarly enriched for CD44^+^CD62L^lo^ effector memory (T_EM_) CD8^+^ T cells in both *Nlrc5^+/+^* and *Nlrc5^−/−^* hosts (Fig. 3B). B16-N-FL TILs, but not B16-N-SA TILs, from *Nlrc5^+/+^* mice also contained increased numbers of CD44^+^CD62L^+^ central memory (T_CM_) CD8^+^ T cells and CD44^lo^CD62L^+^ naive (T_N_) CD8^+^ T cells, whereas these cells were not increased in *Nlrc5^−/−^* hosts (Fig. 3C,D). NLRC5 expression in tumors also augmented the activation status of CD4^+^ T cells and their infiltration into tumors. TILs from B16-N-FL tumors of *Nlrc5^+/+^* mice harbored significantly higher numbers of CD69^+^ T_E_, CD44^+^CD62L^lo^ CD4 T_EM_ and CD44^+^CD62L^+^ CD4 T_CM_ cells, whereas these CD4^+^ T cell subsets were present in elevated numbers in the DLN of B16-N-SA tumors in *Nlrc5^+/+^*hosts (Fig. 3E-G). These cells were not enriched in the TILs of *Nlrc5^−/−^* hosts bearing B16-N-FL or B16-N-SA tumors, except for the elevated number of T_EM_ CD4^+^ T cells in B16-N-SA TILs (Fig. 3E-G). However, the DLN of *Nlrc5^−/−^* hosts bearing B16-N-FL or B16-N-SA tumors harbored increased numbers of all CD4^+^ T cell subsets compared to the DLN of B16-V tumors (Fig. 3E-H). Collectively, these data indicate that NLRC5-FL and NLRC5-SA expressed in tumor cells promote the activation and differentiation of CD8^+^ as well as CD4^+^ T cells in DLN even in the absence of endogenous NLRC5 expression in host APCs, and infiltration of activated and differentiated T cells into tumors.

**Figure 3.**
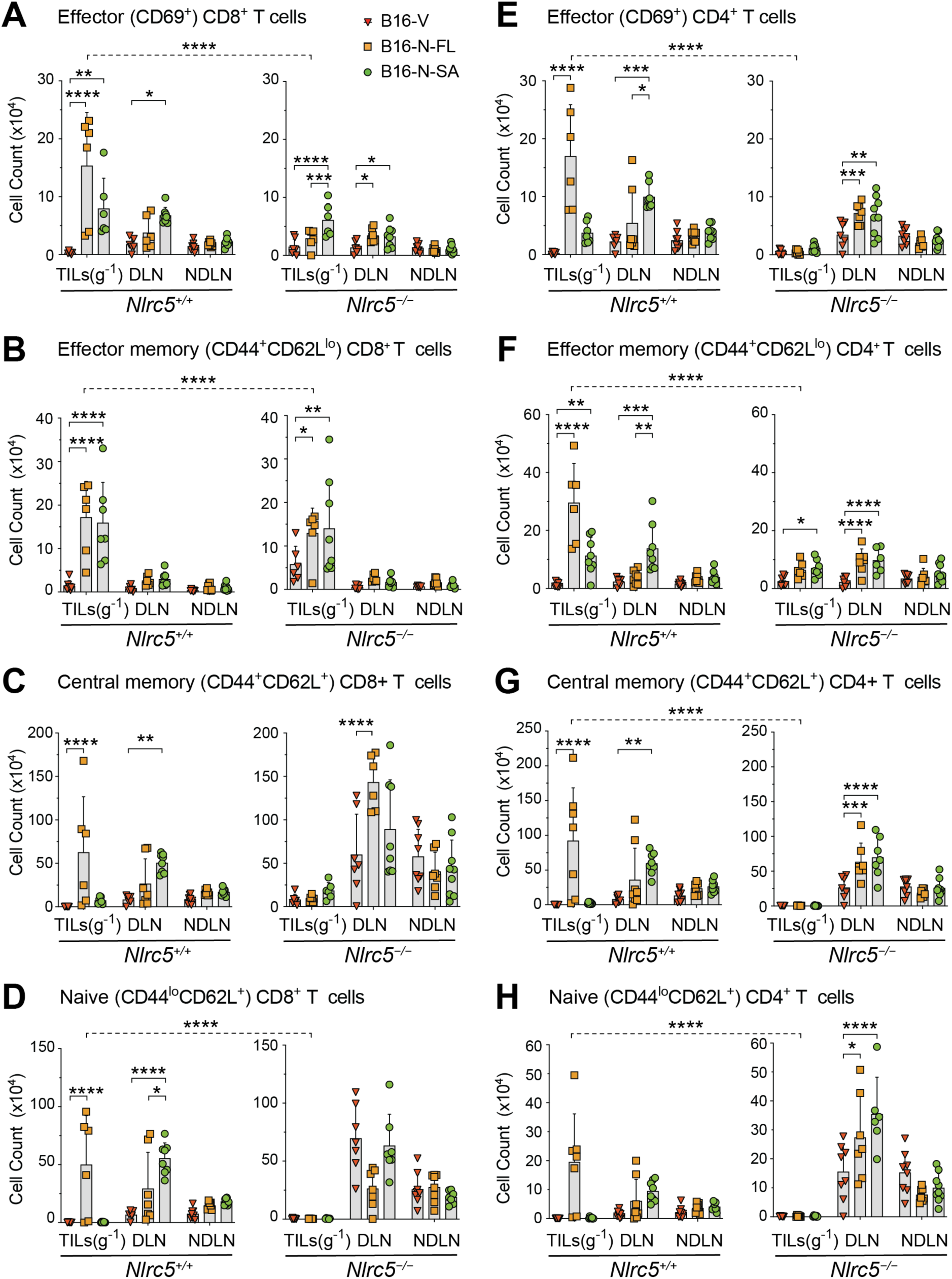
Expression of NLRC5-FL or NLRC5-SA in tumor cells promotes T lymphocyte activation, differentiation and infiltration into tumors in *Nlrc5^+/+^* and *Nlrc5^−/−^* hosts. TILs and cells from DLN and NDLN from B16-V, B16-N-FL or B16-N-SA tumor bearing *Nlrc5^+/+^* and *Nlrc5^−/−^* hosts were stained with fluorochrome conjugated antibodies for activation and differentiation markers of T lymphocytes and analyzed by flow cytometry (Supplementary Fig. S2). Data shown are from 6-9 mice per group from two independent experiments. Absolute numbers of CD8^+^ and CD4^+^ effector (A,E), effector memory (B, F), central memory (C, G) and naïve (D, H) T cells are shown in TILs (g^-1^), DLN and NDLN. Statistics: Mean + SD. Two-way ANOVA with Tukey’s multiple comparison test. * *p* ≤0.05, ** *p* ≤0.01, *** *p* ≤0.001, **** *p* ≤0.0001.

### Tumor cell-intrinsic NLRC5 expression alters the tumor microenvironment

B16-N-FL and B16-N-SA tumors resected from *Nlrc5^+/+^*or *Nlrc5^−/−^* hosts were discernibly firm in consistency compared to B16-V tumors, suggesting an altered tumor microenvironment (TME) in NLRC5 expressing tumors. To characterize the TME, tumor sections were stained with Sirius red to evaluate collagen matrix deposition (Supplementary Fig. S4A) and evaluated for collagen producing myofibroblasts by IF staining of α-smooth muscle actin (αSMA) (Supplementary Fig. S4B). Both B16-N-FL and B16-N-SA tumors showed extensive collagen deposition and accumulation of αSMA^+^ myofibroblasts compared to B16-V tumors, which was confirmed by quantification of positively stained areas (Supplementary Fig. S4C, D).

NLRC5 expression in endothelial cells has been reported to promote angiogenesis ^33^. We observed that B16-N-FL tumors were reddish in color in NSG, *Nlrc5^+/+^* and *Nlrc5^−/−^* hosts compared to the black color of B16-V tumors, whereas B16-N-SA tumors displayed red to brown color (Fig. 1C Supplementary Fig. 1D). As B16-N-FL and B16-N-SA tumors harbor more adaptive and innate immune cells than B16-V tumors, we posited that NLRC5-FL and NLRC5-SA promote tumor vascularization that would facilitate immune cell infiltration. Tumor vascularization can arise from angiogenesis, wherein new capillaries arise from proliferation of endothelial cells (EC), as well as via non-angiogenic mechanisms ^34,35^ such as vasculogenic mimicry (VM), wherein cancer cells acquire EC-like properties and form vessel-like structures ^35–39^. Whereas the expression of CD31 characterizes vascular endothelial cells, tumor cells undergoing VM can be detected using fluorescently labelled tomato lectin (TL) that binds luminal glycocalyx ^40,41^. B16-N-FL and B16-N-SA tumors displayed significantly increased staining for CD31 as well as TL compared to B16-V tumors in both *Nlrc5^+/+^* and *Nlrc5^−/−^* hosts (Fig. 4A-C). CD31 and TL staining showed prominent overlapping in B16-N-FL and B16-N-SA tumors in *Nlrc5^+/+^* mice, but much less in *Nlrc5^−/−^* hosts (Fig. 4A). Moreover, *NLRC5* expression showed a highly significant positive correlation with the expression of *PECAM1* (CD31) in the TCGA data on skin cutaneous melanoma (SKCM) (Fig. 4D).

**Figure 4.**
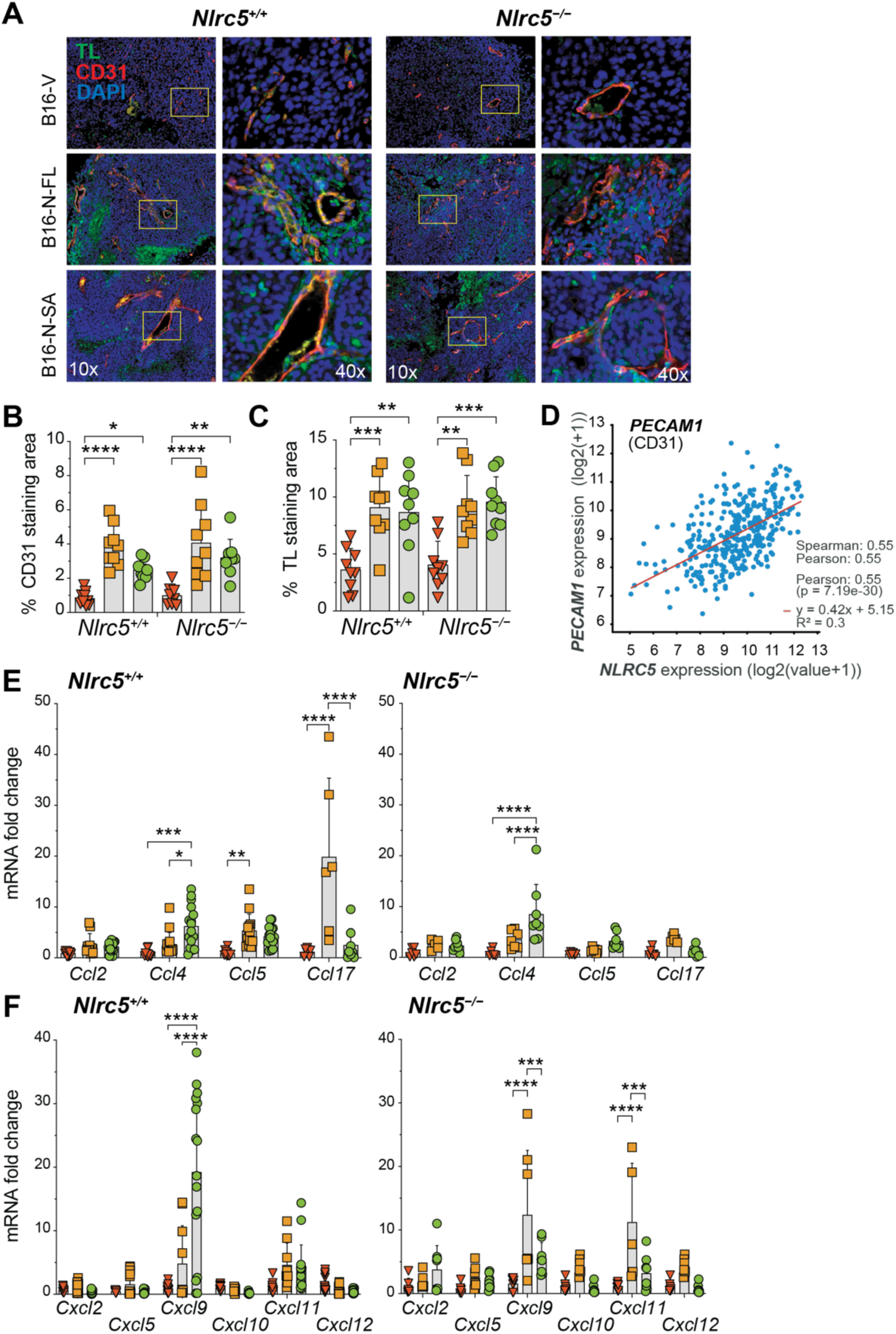
NLRC5-FL or NLRC5-SA expression in tumor cells increases tumor vascularization and modulates chemokine gene expression in *Nlrc5^+/+^* and *Nlrc5^−/−^* hosts. (A) FFPE sections of B16-V, B16-N-FL and B16-N-SA tumors resected from *Nlrc5^+/+^* and *Nlrc5^−/−^* hosts at the tumor endpoint were evaluated for vascular endothelial cells (CD31) and the binding of tomato lectin (TL) by immunofluorescence staining. Representative staining patterns in each tumor are shown at 10× and 40× magnification. (B, C) Percentage of CD31 and TL staining areas was calculated from nine random fields from three tumors per group. (D) Correlation between NLRC5 and PECAM1 (CD31) expression in human skin cutaneous melanoma (SKCM) in the TCGA dataset. (E, F) Total RNA extracted from 6-8 tumors per group were analyzed by RT-qPCR for the indicated CCL. (E) and CXCL (F) chemokine genes. Statistics: Mean + SD. (B,C) Ordinary one-way ANOVA with Tukey’s multiple comparison test. (E,F) Two-way ANOVA with Tukey’s multiple comparison test. * *p* ≤0.05, ** *p* ≤0.01, *** *p* ≤0.001, **** *p* ≤0.0001.

The above findings suggested that increased vascularization driven by NLRC5 expressed in tumor cells could facilitate the trafficking of immune cells into B16-N-FL and B16-N-SA tumors. As immune cell infiltration into tumors is influenced by chemokines in the TME ^42^, we evaluated the expression of CCL and CXCL chemokine genes in tumor tissues. B16-N-FL tumors displayed elevated expression of *Ccl5* in *Nlrc5^+/+^* mice, whereas B16-N-SA tumors showed elevated expression of *Ccl4* compared to B16-V or B16-N-FL tumors in *Nlrc5^+/+^* and *Nlrc5^−/−^* hosts (Fig. 4E). B16-N-FL tumors also showed elevated expression of *Ccl17* compared to B16-V and B16-N-SA tumors in *Nlrc5^+/+^* but not in *Nlrc5^−/−^* mice. Among the CXCL chemokines, *Cxcl9* was elevated in B16-N-SA tumors from *Nlrc5^+/+^* mice, whereas *Cxcl9* and *Cxcl11* were upregulated in B16-N-FL tumors from *Nlrc5^−/−^* mice (Fig. 4E,F).

### High NK cell marker expression in B16-N-FL and B16-N-SA tumors

TILs from B16-N-FL and B16-N-SA tumors or their DLN contained significantly more NK1.1^+^ NK, NKT and iNKT cells than B16-V tumors (Fig. 2E-G). As the activation of these cells can be modulated by several NK cell activating and inhibitory receptors ^29,30,43^, we examined the gene expression of candidate NK cell receptor genes in tumor tissues. Expression of the activating receptor genes *Klrb1c* (NK1.1)*, Klrk1* (NKG2D) and *Klrc2* (NKG2C) were significantly elevated in B16-N-FL and B16-N-SA tumors compared to B16-V tumors in *Nlrc5^+/+^* mice, although the expression of the activating receptor *Ncr1* (NKp46) was not upregulated in NLRC5 expressing tumors (Fig. 5A, upper panel). Notably, the inhibitory receptor *Klrd1* (CD94) also showed upregulation in B16-N-FL and in B16-N-SA tumors from *Nlrc5^+/+^*mice. A similar pattern of activating and inhibitory receptor gene expression also occurred in B16-N-FL tumors in B16-N-SA tumors from *Nlrc5^−/−^* mice (Fig. 5A, lower panel). Expression of *Klra10* (Ly49j), a pseudogene in mouse, and that of the NK cell ligand *Rae1* were not affected in B16-N-FL and B16-N-SA tumors in both *Nlrc5^+/+^* and *Nlrc5^−/−^* mice (Fig. 5A). IFNγ is a key immune effector molecule produced by NK, NKT, iNKT and activated CD8^+^ T cells and CD4^+^ T_H1_ cells. B16-N-SA tumors from *Nlrc5^+/+^* mice showed increased induction of *Ifng* and *Ifnb* genes compared to B16-V tumors (Fig. 5B, upper panel). Upregulation of these genes also occurred in B16-N-FL tumors, but their increase was not statistically significant. On the other hand, B16-N-FL tumors showed significant induction of *Ifng* and *Ifna* genes in *Nlrc5^−/−^* mice (Fig. 5B, lower panel). These observations support the idea that NLRC5 expression in tumors promotes the recruitment of T, NK, NKT and iNKT cells and their effector functions, which could be modulated by endogenous NLRC5 expression in host cells.

**Figure 5.**
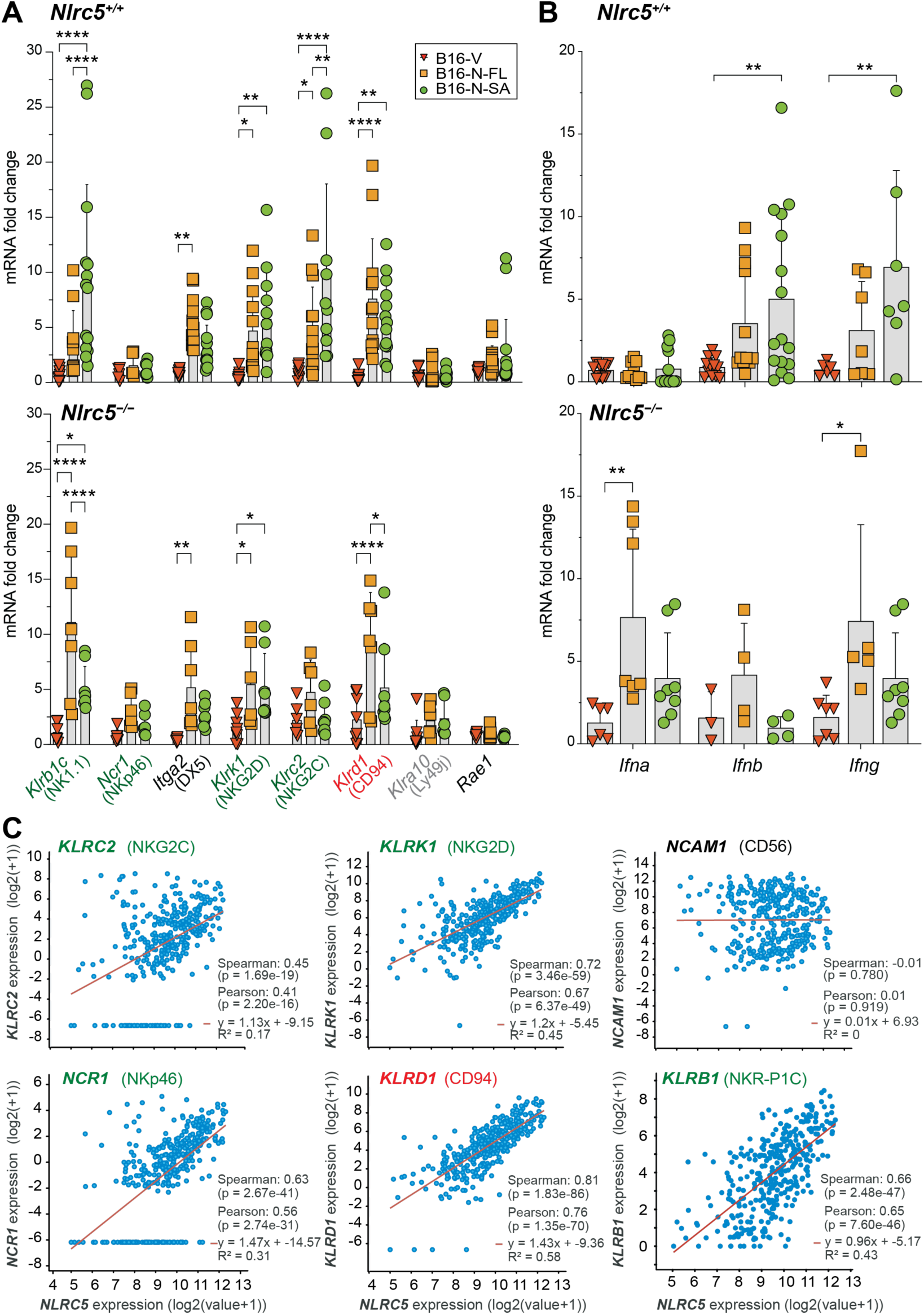
NLRC5 expression in tumor cells upregulates NK cell markers expression in B16 mouse melanoma and in human melanoma. (A) Expression of NK cell marker genes in B16-V, B16-N-FL and B16-N-SA tumors. RNA extracted from tumor tissues resected from *Nlrc5^+/+^* and *Nlrc5^−/−^* hosts (6-8 tumor tissues per group) at the tumor endpoint were evaluated for the expression of indicated NK marker genes by RT-qPCR. Genes indicated in green font are activation markers (*Klrb1c, Ncr1, Klrk1, Klrc2*) with the inhibitory receptor *Klrd1* in red, and the *Itga2* marker and the NK cell ligand *Rae1* in black. *Klra10* is a pseudogene in mouse. Commonly used marker names are given in parenthesis. (B) Gene expression of interferon genes *Ifna*, *Ifnb* and *Ifng* in B16-V, B16-N-FL and B16-N-SA tumor tissues (3-8 tumors per group) resected from *Nlrc5^+/+^* and *Nlrc5^−/−^* hosts. (C) Correlation between the expression of *NLRC5* and NK cell activating (*KLRC2, NCR1, KLRK1, KLRB1*) and inhibitory (*KLRD1*) receptor genes in the human TCGA-SKCM dataset. (A, B) Statistics: Mean + SD. Two-way ANOVA with Tukey’s multiple comparison test. * *p* ≤0.05, ** *p* ≤0.01, **** *p* ≤0.0001.

It has been reported that NLRC5 expression is associated with the recruitment and activation of CD8^+^ T cells but not NK cells in human melanoma, based on the lack of correlation between NLRC5 and the NK cell marker CD56 (NCAM1) ^8^. As our findings show infiltration of NK1.1^+^ NK, NKT, iNKT cells into B16-N-FL and B16-N-SA tumors (Fig. 2), and these tumor tissues express several NK cell activating/inhibitory receptor genes (Fig. 5A), we evaluated the correlation between the expression of *NLRC5* and NK cell receptor genes in the TCGA-SKCM dataset. We found a strong positive correlation between *NLRC5* and several NK activating receptor genes *KLRC2* (NKG2C), *KLRK1* (NKG2D), *NCR1* (NKp46) and *KLRB1* (NKR-P1C, the human equivalent of NK1.1) as well as the inhibitory receptor *KLRD1* (CD94), even though no correlation was observed between *NLRC5* and the NK cell marker *NCAM1* (CD56) expression (Fig. 5C). *NLRC5* expression significantly correlated with *CD8A* and *CD4*, and with transcription factors *TBX21* (TBET), *EOMES*, *GATA3*, *RORC* (RORγT), which are implicated in the functional differentiation of CD4 and CD8 T cells, NK and ILC cells (Supplementary Fig. S5A-B). Accordingly, *NLRC5* expression strongly correlated with *IFNG* expression, but minimally with *IFNB1* or *IL22* (Supplementary Fig. S5C). Moreover, *NLRC5* expression correlated with the classical MHC-Ia genes *HLA-A, B* and *C* as well as with the non-classical MHC-Ib genes *HLA-E, F* and *G*, and MHC-II genes *HLA-DRA, DRB1* and *DPA1* (Supplementary Fig. 6A-C). These data support the possibility that NLRC5 expression in tumor cells not only promotes the activation of CD8^+^ T cells but also NK cells and other innate and adaptive immune cells that produce IFNγ to mediate antitumor functions.

### Control of B16-N-FL and B16-N-SA tumors requires both CD8^+^ T and NK1.1^+^ cells

B16-N-FL and B16-N-SA tumors harbored elevated numbers of NK1.1^+^ (NK, NKT and iNKT) cells in TILs or DLN (Fig. 2E-G). Therefore, we investigated whether NK1.1^+^ cells are also needed for tumor control, in addition to CD8^+^ T cells. Depletion of either CD8^+^ T cells or NK1.1^+^ cells resulted in a significant increase in the growth of B16-N-FL and B16-N-SA tumors, which were comparable to the growth of B16-V tumors, in *Nlrc5^+/+^* hosts (Fig. 6A-C), indicating that both CD8^+^ T cells and NK1.1^+^ cells play crucial roles in controlling NLRC5-expressing tumors. CD8^+^ T cell depletion increased the growth of B16-N-FL and B16-N-SA tumors in *Nlrc5^−/−^* mice as well (Fig. 6D, F). However, depletion of NK1.1^+^ cells increased the growth of B16-N-SA but not B16-N-FL tumors in *Nlrc5^−/−^* mice (Fig. 6D-F).

**Figure 6.**
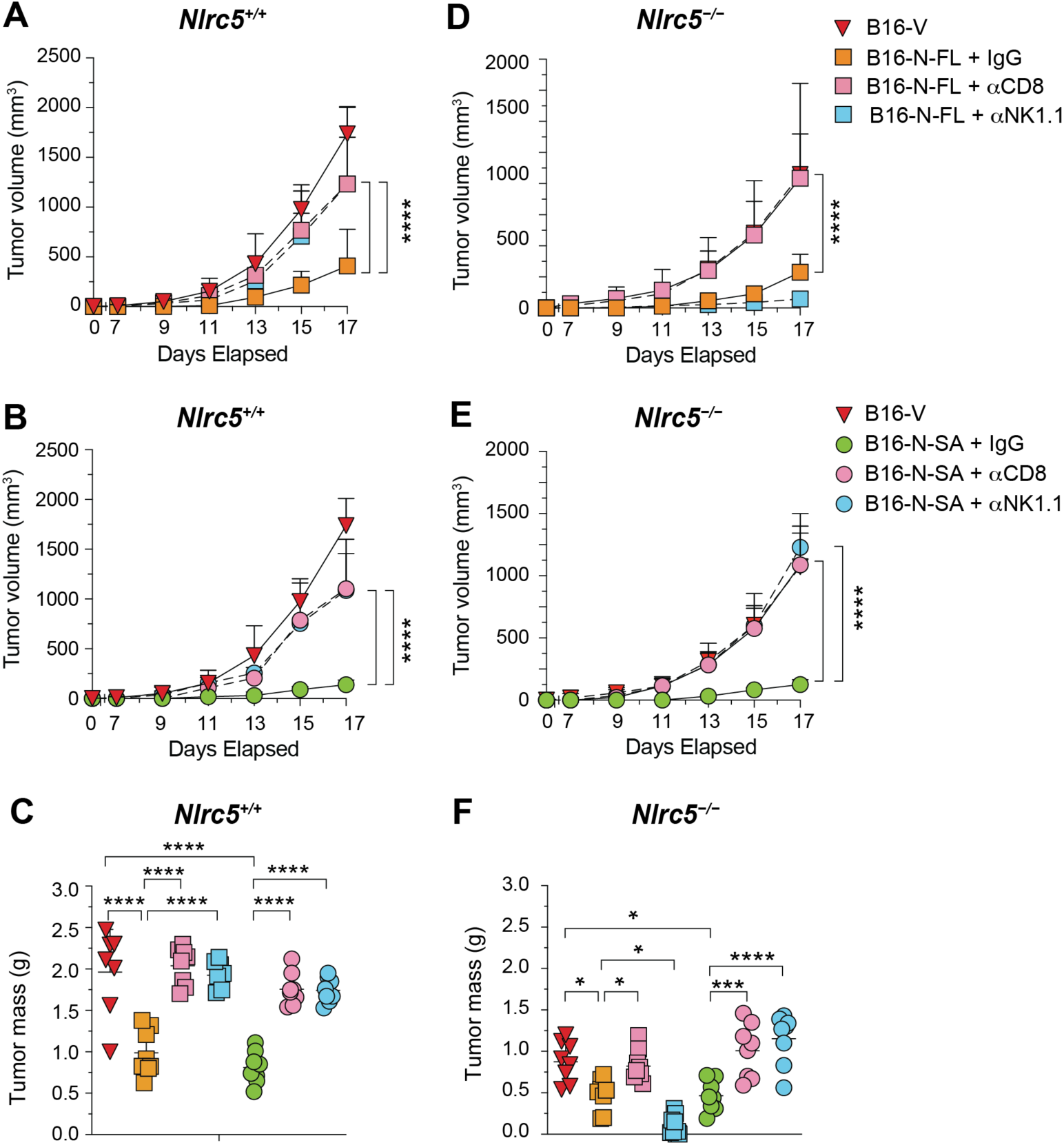
Control B16-N-FL and B16-N-SA tumors requires both CD8^+^ T and NK1.1^+^ cells. (A) *Nlrc5^+/+^* (A-C) and *Nlrc5^−/−^* (D-F) mice were injected with anti-CD8α, anti-NK1.1 (PK136) or control (IgG2a) antibodies to deplete CD8+ or NK1.1+ cells 24 h prior to and 24 h following subcutaneous implantation of 2 × 10^5^ B16-N-FL (A,C,D) or B16-N-SA (B,E,F) cells. Additional doses of Ab were injected 7 and 14 days later. Mice injected with B16-V cells served as additional controls for tumor growth comparison. Data from eight mice per group from two different experiments. (A-C) Growth kinetics (A,B) and tumor weight at sacrifice (C) for B16-N-FL and B16-N-SA tumors in *Nlrc5^+/+^*hosts. (D-F) Growth kinetics (D,E) and tumor weight at sacrifice (F) for B16-N-FL and B16-N-SA tumors in *Nlrc5^−/−^*hosts. Statistics: Mean + SD. Two-way ANOVA with Tukey’s multiple comparison test. * p≤0.05, *** p≤0.001, **** p≤0.0001.

### B16-N-FL and B16-N-SA cells downmodulate lysosome and phagosome pathways

To understand the molecular mechanisms that contribute to the efficient control of B16-N-FL and B16-N-SA tumors compared to B16-V tumors, we carried out total proteome analysis of B16-V, B16-N-FL and B16-N-SA cells. Quality control data on the proteome profiles of quadruplicate samples from each cell line are shown in Supplementary Fig. S7. Nearly 6000 proteins were detected in each sample with a combined total of 6208 individual proteins, and missing value pattern analysis showed a random distribution of missing and detected proteins (Supplementary Fig. S7A, B). Principal component analysis of the proteomic data showed clear separation of quadruplicate samples from each cell line and a good correlation among the replicates (Supplementary Fig. S7C,D). Volcano plot of proteins detected in B16-N-FL and B16-V cells showed significant modulation of 1372 differentially expressed proteins (DEPs), with 583 upregulated and 789 downregulated proteins, whereas B16-N-SA cells showed 862 DEPs compared to B16-V cells with 412 upregulated and 450 downregulated proteins (Fig. 7A, B). Notably, PMEL, TYR, DCT and TYRP1 proteins, which carry dominant antigenic epitopes recognized by CTLs ^44^, were downmodulated in B16-N-FL cells, and the latter three in B16-N-SA cells, compared to B16-V cells (Fig. 7A,B). A comparison of B16-N-SA and B16-N-FL cells revealed 1200 DEPs with 740 upregulated and 460 downregulated proteins, with relative upregulation of PMEL, TYRP1 and DCT in B16-N-SA cells compared to B16-N-FL cells (Fig. 7C). Assessing the degree of overlap between proteins significantly upregulated or downregulated in B16-N-FL compared to B16-V cells, B16-N-SA compared to B16-V cells, and B16-N-SA compared to B16-N-FL cells showed that B16-N-FL cells contained more unique proteins than B16-N-SA cells, which also harbored a significant number of uniquely modulated proteins (Fig. 7D).

**Figure 7.**
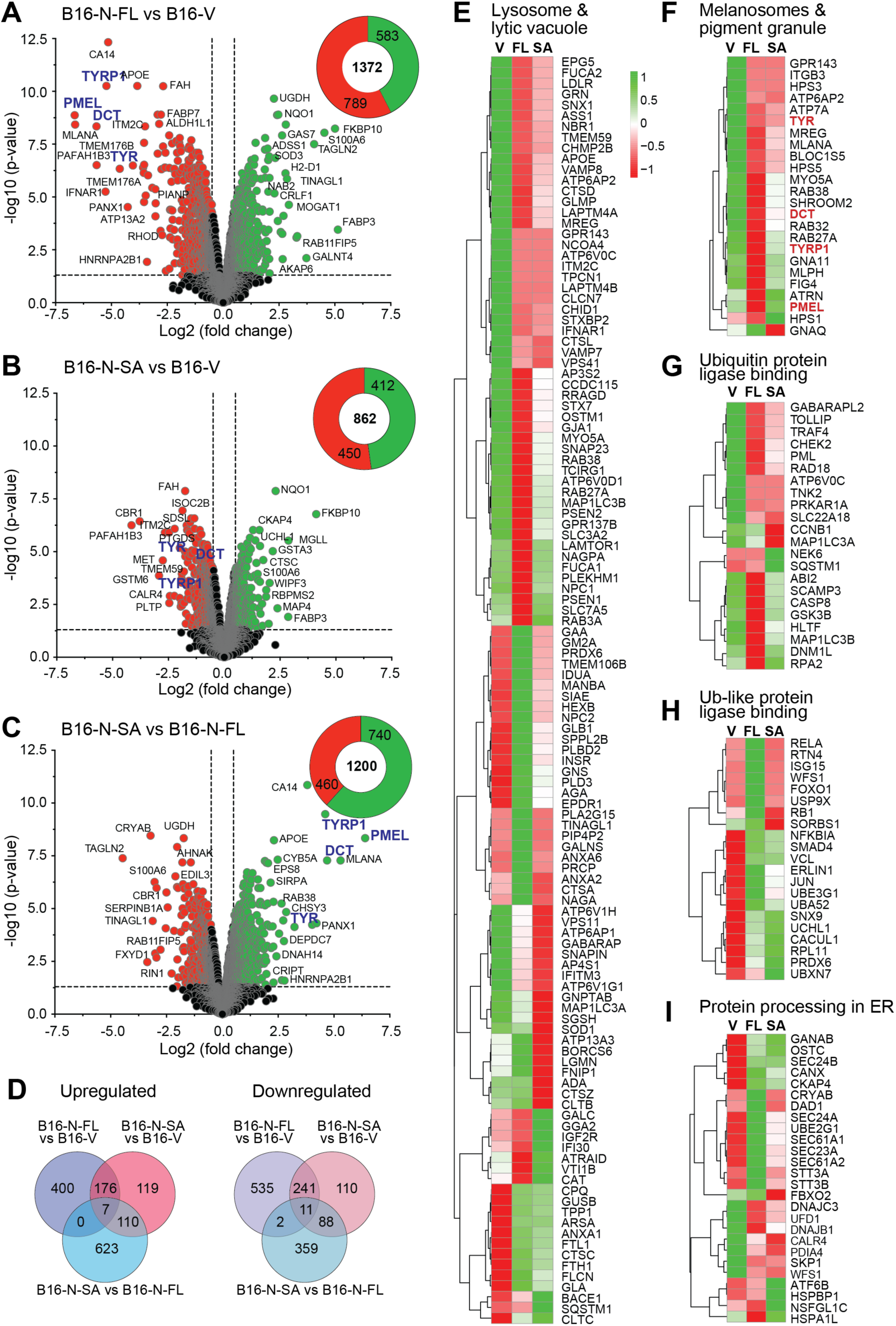
Proteomes of NLRC5-FL and NLRC5-SA display overlapping and distinct modulation of cellular proteomes. Quadruplicate samples of proteins extracted from B16-V, B16-N-FL and B16-N-SA cells were analyzed by mass spectrometry. (A-C) Volcano plots of proteins detected in B16-N-FL (A) and B16-N-SA (B) cells compared to B16-V cells, and in B16-N-SA cells compared to B16-N-FL cells (C). Upregulated proteins are indicated in green color and downregulated proteins in red color. Number of upregulated, downregulated and total DEPs are shown in the pie chart. (D) Venn diagrams showing upregulated and downregulated proteins in the comparisons shown in A-C. (E-I) Heatmap analyses of proteins in GO:CC terms lysosome and lytic vacuole (E) and melanosome and pigment granule (F), GO:MF terms ubiquitin protein ligase binding (G) and ubiquitin-like protein ligase binding (H) and the pathway term protein processing in the ER (I) in B16-V, B16-N-FL and B16-N-SA cells. Normalised abundance values (calculated using complete-linkage clustering and Euclidean distance method) are color coded on a scale with green and red colors representing upregulation and downregulation, respectively.

We also observed enrichment of the GO:cellular compartments (CC) terms ‘lytic vacuole’ and ‘lysosome’ among the upregulated proteins in B16-N-FL and B16-N-SA cells (Supplementary Fig. S8A,B). Interestingly, these two GO terms were also enriched among the downmodulated proteins in B16-N-FL cells (Supplementary Fig. S8D) Lysosome was also enriched in the pathway analysis of upregulated proteins in B16-N-FL and B16-N-SA cells (Supplementary Fig. S9). Heatmap of DEPs modulated within the lysosome and lytic vacuole GO:CC terms revealed a complex pattern with nearly half of the DEPs downregulated in B16-N-FL cells and to a lesser extent in B16-N-SA cells, although several other proteins were downregulated to a greater extent in B16-N-SA than in B16-N-FL cells (Fig. 7E). About a third of DEPs showed low expression in B16-V cells, of which many were upregulated only in B16-N-FL and certain others only in B16-N-SA cells (Fig. 7E). The GO:CC terms ‘melanosome’ and ‘pigment granule’ were significantly enriched among the downregulated proteins in B16-N-FL and B16-N-SA cells (Supplementary Fig. S8D,E), and many melanoma proteins harboring CTL epitopes (PMEL, TYR, DCT and TYRP1) are downmodulated in these cells (Fig. 7A,B). A heatmap analysis of DEPS within the GO terms melanosome and pigment granule showed marked downregulation of most DEPs including PMEL, TYR, DCT and TYRP1 in B16-N-FL cells and to a lesser extent or remained unmodulated in B16-N-SA cells (Fig. 7F).

The GO terms of molecular functions (MF) term ‘ubiquitin protein ligase binding’ and ‘ubiquitin-like protein ligase binding’ were enriched in B16-N-FL cells but not in B16-N-SA cells (Supplementary Fig. S8A,B). As ubiquitinated proteins are targeted to proteasomes to generate peptides for MHC-I binding, we examined the expression patterns of DEPs in these two GO:MF terms. DEPs involved in ubiquitin protein ligase binding were mostly downregulated in B16-N-FL cells whereas DEPs involved in ubiquitin-like protein ligase binding were upregulated (Fig. 7G,H). On the other hand, only half of the DEPs in ubiquitin protein ligase binding were downregulated, and half of the DEPs in ubiquitin-like protein ligase binding are upregulated in B16-N-SA cells. The molecular function term ‘protein processing in the ER’, enriched among the upregulated proteins in B16-N-FL cells (Supplementary Fig. S9), contained several upregulated and downregulated proteins, whereas B16-SA cells showed fewer and lesser changes (Fig. 7I).

Next, we focussed on DEPs that are shared between B16-N-FL and B16-N-SA cells compared to B16-V cells (Fig. 8A). Among the 465 shared proteins, the GO terms lytic vacuole, lysosome, melanosome and pigment granule were highly enriched in B16-N-FL and B16-N-SA cells (Fig. 8B). Pathway analysis showed significant enrichment of phagosomes and autophagy in B16-N-FL and B16-N-SA cells (Fig. 8C). Heatmap analysis of DEPs in the phagosome pathway prominently showed downmodulation of the V-type ATPase subunits ATP6V1B2 and TP6V1E1 to a greater extent in B16-N-FL cells, whereas ATP6V1G1 ATP6V1A, ATP6V1F, ATP6V1G1 and ATP6V1D1 were downmodulated to a greater extent in B16-N-SA cells (Fig. 8D). It is noteworthy that ‘proton-transporting V-type ATPase complex’ was enriched among the GO:CC terms (Fig. 8B). Most DEPs related to the autophagy pathway were downmodulated in B16-N-FL and B16-N-SA cells, whereas some were upregulated, and SQSTM1 (p62) was differentially expressed in B16-N-FL and B16-N-SA cells (Fig. 8E). Notable among the downregulated proteins are NBR1, MAP1LC3B (LC3B) and GABARAPL2, which are implicated in the degradation of MHC-I in autophagosomes ^45,46^. As autophagy in tumor cells promotes cross presentation of tumor antigens ^47^, our findings suggest that NLRC5 expressed in cancer cells modulates vacuolar compartments that could impact tumor antigen presentation by APCs in DLN.

**Figure 8.**
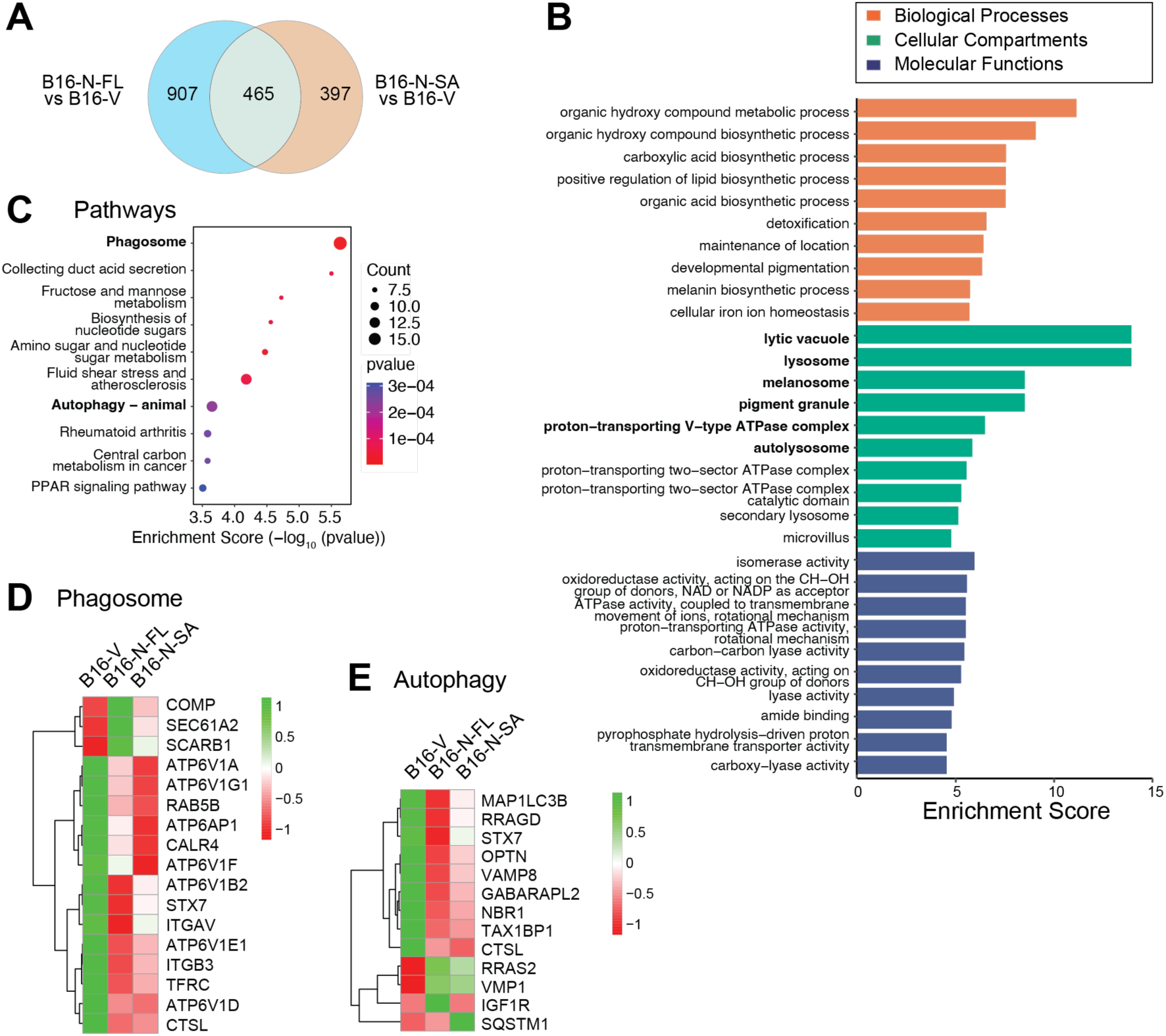
Enrichment of phagosome and autophagy pathway proteins in the shared proteome of NLRC5-FL and NLRC5-SA cells. (A) Venn diagram showing shared and unique proteins differentially expressed in B16-N-FL and B16-N-SA cells compared to B16-V cells. (B) GO analysis of DEPs shared between B16-N-FL and B16-N-SA cells. (C) Pathway analysis of DEPs shared between B16-N-FL and B16-N-SA cells. Key enrichment terms in B and C are indicated in bold font. (D,E) Heatmap analyses of phagosome and autophagy pathway proteins differentially expressed in B16-N-FL and B16-N-SA cells compared to B16-V cells.

## Discussion

Downmodulation of NLRC5 in many cancers and its correlation with low MHC-I expression and poor response to immune checkpoint therapy have raised the prospects of exploiting NLRC5 to restore tumor immunogenicity in MHC-I-low immune evasive cancers ^6,8,11^. We have previously reported that expression of NLRC5-FL or NLRC5-SA in the preclinical models of B16 melanoma and EL4 thymoma promotes tumor control ^10,15^. We have recently shown using *Nlrc5^-/-^*mice that NLRC5 is required for cancer immune surveillance and cancer immunoediting ^13^. In the present work, we dissected the role of NLRC5 expressed in tumor cells from that of its expression in APCs and other cells of the host. Our findings reveal that NLRC5 expressed in tumor cells promote activation of CD8^+^ T cells as well as NK1.1+ cells and CD4^+^ T cells and their infiltration into tumors, and that these processes occur even in the absence of NLRC5 expression in host APCs. Our findings also indicate that NLRC5 expression in tumor cells modulates the TME that supports immune cell infiltration and tumor control, and that NLRC5 expression in other host cells impacts NK cell-dependent tumor control.

NLRC5-deficient mice have been reported to display impaired CTL activation against intracellular pathogens and viruses ^25,26,48^. Mice lacking NLRC5 in dendritic cells (DC) showed reduced MHC-I-restricted antigen specific CD8^+^ T cell responses in the intestine following oral rotavirus infection ^48^, suggesting a role for NLRC5 in antigen cross presentation by APCs. Tumor-reactive CD8^+^ T cells are initially activated by tumor antigens in DLN by type 1 conventional DCs (cDC1) via antigen cross-presentation, which can also be mediated by tumor-associated macrophages ^49–51^. Hence, we anticipated that the cross-presentation of tumor antigens by APCs would be less efficient in *Nlrc5^−/−^* mice bearing B16-N-FL or B16-N-SA tumors. In contrast to this prediction, B16-N-FL or B16-N-SA tumors were controlled in *Nlrc5^−/−^* mice as efficiently as, and even better than, in *Nlrc5^+/+^* hosts. These results indicate that NLRC5 expression in APCs is dispensable for tumor antigen cross-presentation to CD8^+^ T cells and suggest that NLRC5 expression within tumor cells may facilitate tumor antigen presentation by host APCs through other mechanisms. Recent reports have shown that tumor antigen presentation via cross-dressing of tumor cell-derived MHC-I:peptide complexes, acquired by cDC1 cells via trogocytosis of tumor cells, is a key pathway to prime naïve CD8^+^ T cells and elicit antitumor immunity ^21,22,52^. Hence, our findings raise the possibility that the loss of NLRC5 expression in tumor cells could compromise tumor antigen presentation via cross-dressing, thereby hindering the iterative processes in the cancer immunity cycle ^53^. Our data also suggest that improved antigen presentation by NLRC5 in tumor cells may also widen the repertoire of tumor antigens presented to CD8^+^ T cells. Despite showing significant downmodulation of major tumor antigens such as PMEL1, DCT, TYR and TRP1 bearing dominant CTL epitopes ^54^, B16-N-FL and B16-N-SA tumors displayed increased CD8^+^ T cell activation in DLN and their infiltration into tumors. This could arise from more efficient antigen processing resulting from the upregulation of antigen processing machinery within tumor cells expressing NLRC5 as well as from improved transfer of MHC-I:peptide complexes to APCs by cross-dressing, as discussed above. This would result in efficient tumor cell killing and consequent iteration of the tumor immunity cycle. This process could also underlie the marked activation of CD4^+^ T cells in the DLN of mice bearing B16-N-FL and B16-N-SA tumors. Hence, restoring a functional NLRC5 in tumor cells using NLRC5-SA would be a promising approach not only to restore MHC-I but also to facilitate tumor antigen presentation and to reinstate antitumor immunity.

Proteome analysis of B16-N-FL or B16-N-SA cells provides some clues to potential mechanisms by which NLRC5 expressed in cancer cells could impact tumor antigen presentation by APCs. Differentially expressed proteins shared between B16-N-FL or B16-N-SA cells showed enrichment of proteins residing in vacuolar compartments such as lytic vacuole, lysosome and autolysosome, and participate in phagosome and autophagy pathways. Autophagy in cancer cells can promote MHC-I dependent antitumor immune response by promoting cross-presentation of tumor antigens or dampen these responses by promoting MHC-I degradation ^45,47,55^. Variables that determine the differential impact of autophagy on MHC-I dependent antitumor immunity are not yet well understood. As autophagy-low tumours show elevated MHC-I expression and tumour-infiltrating CD8^+^ T cells ^45^, our observations suggest that low NLRC5 expression could be a key determinant of autophagy-driven MHC-I degradation in cancers. In support of this notion, autophagy induction in B16 melanoma cells has been reported to reduce MHC-I expression and that this is overcome by IFNγ treatment ^56^. As NLRC5 is strongly induced by IFNγ and B16 cells are responsive to IFNγ ^32,57^, our findings suggest that NLRC5 induced by IFNγ overcomes autophagy mediated MHC-I downregulation by attenuating the autophagic process. Thus, delivering NLRC5-SA would not only promote MHC-I-dependent antitumor immune responses but also target autophagy and its pro-tumorigenic effects ^55,58^.

The initial study on NLRC5 expression across cancers in TCGA-SKCM transcriptomic data showed that the mRNA levels of *NLRC5* positively correlated with those of classical MHC-Ia molecules (*HLA-A, HLA-C, B2M*), MHC-I antigen presentation pathway genes (LMP2/*PSMB9*, LMP8/*PSMB8*, *TAP1*), CD8A and effector molecules of cytotoxic T cells (*GZMA, PRF1*), but not with the NK cell marker CD56 (*NCAM1*), suggesting that NLRC5 may not impact NK cell recruitment and their antitumor functions ^8^. However, we observed significant enrichment of NK, NKT and iNKT cells in TILs of B16-N-FL tumors in *Nlrc5^+/+^* mice and in TILs of B16-N-SA tumors in *Nlrc5^−/−^* hosts, indicating a role for NLRC5 in promoting innate antitumor immune responses. This notion was supported by elevated mRNA levels of NK cell activating receptor molecules in B16-N-FL and B16-N-SA tumors, and a significant positive correlation between the transcript levels of NLRC5 and NK receptor genes in TCGA-SKCM data, and the requirement of NK1.1^+^ cells for tumor control. However, depletion of NK1.1^+^ cells resulted in increased growth of B16-N-FL and B16-N-SA tumors in *Nlrc5^+/+^* mice but only that of B16-N-FL tumors in *Nlrc5^−/−^* hosts, suggesting that the impact of NLRC5 on antitumor functions of NK1.1^+^ cells could be influenced by NLRC5 expression in other host cells and involves the C-terminal end of NLRC5. Some NK cell receptors recognize non-classical MHC-Ib molecules as ligands, which are implicated in NK cell education and function ^59^. NLRC5 has been reported to transactivate non-classical MHC-Ib molecules H2M3, H2Qa1 and Tla in mice ^25–28^. We also observed a significant positive correlation between the expression levels of NLRC5 and human MHC-Ib molecules HLA-E, HLA-F and HLA-G in TCGA-SKCM (Supplementary Fig. S6). As classical MHC-Ia molecules upregulated by N-FL and N-SA would be inhibitory to NK cells, whether and how NLRC5-induced MHC-Ib contributes to the control of B16-N-FL and B16-N-SA tumors by NK1.1^+^ cells remains to be elucidated.

A notable feature of B16 tumors expressing NLRC5-FL or NLRC5-SA was increased vascularization and collagen deposition. NLRC5 has been reported to promote vascular remodeling in inflammatory settings by attenuating vascular smooth muscle cell proliferation via interaction with PPARγ and by inducing its target genes ^60,61^. NLRC5 promotes angiogenesis in ischemic tissues by increasing endothelial cell (EC) proliferation and reduced B16 tumor growth in EC-specific *Nlrc5^-/-^*mice, suggesting that NLRC5-mediated angiogenesis could promote tumor growth ^33^. This possibility is unlikely because NLRC5 expression in human melanoma positively correlates with CD8^+^ T cell infiltration in tumors and patient survival ^8^. Moreover, we observed that B16-N-FL and B16-N-SA tumors promoted tumor vascularization in both *Nlrc5^+/+^* and *Nlrc5^-/-^*hosts, accompanied by increased immune cell infiltration into tumors despite increased collagen deposition in the TME. Our data indicate that NLRC5-mediated tumor vascularization could result from both neo-angiogenesis and non-angiogenic mechanisms ^35,38^. The occurrence of tumor NLRC5-driven neo-angiogenesis even in *Nlrc5^-/-^* hosts suggests that tumor-derived NLRC5 somehow gains access to host endothelial cells, possibly via tumor-derived exosomes delivering NLRC5 mRNA and protein. Mechanisms underlying this process and NLRC5-driven non-angiogenic tumor vascularization remain to be elucidated. Transcriptional signatures of tumor ECs markedly vary from those of ECs in physiological angiogenesis ^35^. Alterations in tumor ECs not only cause stromal remodeling to aid vessel growth but also establish an immune-hostile TME characterized by impaired trans-endothelial leukocyte migration, diminished proinflammatory signaling, recruitment of immunosuppressive cell types and upregulation of FASL to induce apoptosis of CTLs. Vessel normalization and endothelial reprogramming are some of the approaches aimed at disrupting the immune-suppressive EC-to-immune cell crosstalk to re-establish an immune-supportive TME ^62–66^. Increased infiltration of innate and adaptive antitumor immune cells in B16-N-FL and B16-N-SA tumors indicate that restoring NLRC5 expression would not only reverse MHC-I defects and improve antigen presentation, but also would help establish an immune-supportive TME to facilitate effective tumor control.

One of the mechanisms by which restoring NLRC5 expression in tumors help establish an immune-supportive TME process could be an altered intratumoral chemokine milieu ^42^. CCL4 and CXCL9 chemokine genes upregulated in B16-N-SA tumors are implicated in the recruitment of cDC1 cells and NK and T effector cells, respectively ^67–69^. Suppression of CCL4 production and reduced cDC1 migration into melanomas has been linked to the activation of β-catenin and induction of the transcriptional repressor ATF3 ^67^. Our findings suggest that NLRC5-SA can be exploited to overcome this repression and promote cDC1 recruitment into tumors. Tumor infiltrating cDC1 cells are one of the major producers of CXCL9 that promote recruitment of effector CD8^+^ T cells into the TME ^70^. Even though cDC1 cells also produce CXCL10, another chemokine implicated in recruiting effector T cells ^71^, the *Cxcl10* gene was not upregulated in B16-N-SA tumors. CXCL9, CXCL10 and CXCL11 form a triad of ligands that bind the CXCR3 receptor, which is highly expressed in Th1 cells, CTLs, NK cells and NKT cells, promoting their recruitment to the TME ^72^. IFNγ is strong inducer of CXCL9, CXCL10 and CXCL11 genes, whereas type-I IFNs have differential impact on these genes ^72^. While it is possible that abundant production of IFNγ, presumably from recruited NK cell subsets and effector CD8^+^ T cells, in B16-N-SA tumors of *Nlrc5^+/+^* hosts likely underlies elevated *Cxcl9* gene induction, negligible induction of *Cxcl10* and modest upregulation of *Cxcl11* genes suggest a complex interplay of cellular and soluble mediators impacting the highly variable chemokine milieu of B16-N-FL and B16-N-SA tumors in *Nlrc5^+/+^* and *Nlrc5^-/-^* hosts.

A limitation of our study is that immune cell profiling of TILs and DLN and gene expression studies in B16-N-FL and B16-N-SA tumors were carried out when control B16-V tumors reached the endpoint in *Nlrc5^+/+^* hosts. It is likely immune cell activation in mice bearing B16-N-FL or B16-N-SA tumors occurred much earlier than in mice bearing B16-V tumors, and at different rates in *Nlrc5^+/+^* and *Nlrc5^-/-^*hosts. Studying the immune parameters in tumors, TILs and DLN at earlier time points in mice bearing B16-N-Fl or B16-N-SA tumors will inform how NLRC5-FL and NLRC5-SA impact the kinetics of immune cell activation, their infiltration into tumors and the establishment of antitumor immunity. Another limitation of our study is the focus on lymphoid cells, driven by the requirement of NLRC5 for the transcriptional activation of MHC-Ia and MHC-Ib genes. Characterization of the myeloid cell compartment in TILs and DLN in B16-N-FL or B16-N-SA tumors when control B16-V tumors attain the exponential growth phase would shed light on the dynamics of APCs modulated by NLRC5 expression in tumor cells.

Overall, our findings reveal the tumor cell-intrinsic roles of NLRC5 in mediating tumor control by CD8^+^ T cells and NK1.1^+^ cells, via mechanisms that involve efficient activation of these cells in DLN and their recruitment into tumors by establishing an immune-supportive TME. Despite the noticeable quantitative and qualitative differences between B16-N-FL and B16-N-SA tumors in immune cell recruitment and expression of molecules that can impact immune cell recruitment and activation, their shared proteomes shed light on the potential mechanisms by which NLRC5 can impact diverse aspects of the antitumor immune response. Our study paves a strong foundation for developing NLRC5-SA based cancer immunotherapeutic approaches that can complement other ongoing efforts to reinstate antitumor immunity in immune escape cancers.

## Materials and Methods

### Cell culture

B16F10 murine melanoma cells were obtained from American Type Culture Collection (ATCC, Virginia, USA) and cultured in 5% Fetal Bovine Serum (FBS) containing GibcoTM DMEM (Gibco, #11995065) supplemented with 100U/ml Penicillin/Streptomycin (#P4333), 1mM Sodium pyruvate (#600-11-EL) and 10mM HEPES (#H0887 Sigma-Aldrich). Plasmid vectors expressing full-length human NLRC5 or the engineered version called NLRC5 super-activator (NLRC5-SA) have been previously described ^15^. NLRC5-SA, comprised of the N-terminal domain of NLRC5 and the death domain and LRRs of NLRA (CIITA), induces MHC-I expression as efficiency as NLRC5-FL ^17^. We have previously reported B16-F10 cells stably expressing the control Vector (B16-V), NLRC5-FL (B16-NLRC5-FL) or NLRC5-SA (B16-NLRC5-SA) ^10,15^. The growth rate of the three cell lines was compared using the WST-8 (water-soluble Tetrazolium-8: 2-(2-methoxy-4-nitrophenyl)-3-(4-nitrophenyl)- 5-(2,4-disulfophenyl)-*2H*-tetrazolium) assay kit (CCK-8 Dojindo Molecular Technologies, #CK04) following manufacturer’s instructions. Briefly 5000 cells in medium were seeded per well in a 96 cell well plate in triplicates and 100μL WST-8 reagent was added 24, 48 or 72 h later and incubated for 2 h. Color development, indicative of cell number and viability, was measured at 450 nm wavelength using the SPECTROstar^Nano^ (BMG Labtech, Germany) spectrometer.

### Mice and husbandry

NOD.*scid.gamma* (NSG) mice (NOD.Cg-*Prkdc^scid^ Il2rg^tm1Wjl^*/SzJ) were purchased from the Jackson Laboratory (Bar Harbor, ME USA). *Nlrc5^−/−^*and *Nlrc5^+/+^* control mice in the C57BL/6J background have been previously described ^73^. Briefly, *Nlrc5^−/−^*mice were generated by crossing *Nlr5*-floxed mice with CMV-*cre* deleters obtained from Dr. Dana Philpott ^48^. However, loss of the *Nlr5^fl/fl^* parental stock during the COVID-19 pandemic necessitated crossing *Nlrc5^−/−^* mice with C57BL/6 (Jax Strain #: 000664) mice to generate *Nlrc5^+/+^* littermate controls, following previously reported genotyping protocol^48^. Functional impact of NLRC5 deficiency was confirmed by the impaired upregulation of MHC-I expression, and MHC-I, *B2m* and APM genes in splenocytes following IFNγ stimulation ^13^. All strains of mice were housed in the same breeding and experimentation rooms of the Université de Sherbrooke specific pathogen-free animal facility. Mice were housed in ventilated cages with a 14/10 h day/night cycle and fed with normal chow *ad libitum*. All experiments on mice were performed during daytime with the approval of the Université de Sherbrooke Ethics Committee for Animal Care and Use (Protocol ID 2023-4043).

### Tumor growth and tissue processing

To evaluate tumor formation by B16-V, B16-NLRC5-FL and B16-NLRC5-SA cell lines, 2 × 10^5^ cells suspended in 50 µL phosphate-buffered saline (PBS) were subcutaneously injected into the right flanks of 8 weeks old NSG, *Nlrc5^−/−^* and *Nlrc5^+/+^* mice. The mice were monitored daily, and upon palpable tumor formation, tumor dimensions (length and width) were recorded every two days using a digital vernier caliper. Tumor volume was calculated using the ellipsoid formula: V = ½ x Length x Width^2^. At the endpoint (2 cm in diameter in any one direction), mice were euthanized and tumor mass excised and photographed at a constant zoom factor of 1.7 in a Petri dish. The tumors were sectioned into small fragments. Portions of the tumors were snap-frozen for protein analysis and preserved in RNAprotect Tissue Reagent (Qiagen, #76106) for gene expression studies. For histology tissue pieces were fixed in 4% paraformaldehyde (PFA Sigma-Aldrich, #P6148) overnight, rinsed with 70% ethanol, placed in cassettes and processed for paraffin embedding. Fresh tissues were used for flow cytometry analysis.

### Histology and immunohistochemistry

Formalin-fixed paraffin-embedded (FFPE) tumor sections (4-5 µm) were deparaffinized, rehydrated, stained with hematoxylin and counterstained with eosin (H&E). After mounting with Permount^TM^ mounting medium (#SP15-100

Fisher Scientific) and drying, the slides were scanned using the Digital NanoZoomer (Hamamatsu Photonics 2.0-RS). For staining the collagen matrix, deparaffinized tissue sections were immersed in 0.05% fast green (Sigma, #F7252) for approximately 4 min and then in acidified water (1% acetic acid) for 1 min. After washing with water for 5 min, slides were immersed in picrosirius red staining solution (1 g of Direct Red 80 [Sigma-Aldrich, #365548] in 500 mL saturated picric acid) at room temperature for 1 h. The staining intensity was enhanced using acidified water. Slides were dehydrated, mounted and then scanned using the Digital NanoZoomer. Images were analyzed using the Nanozoomer Digital Pathology (NDP View 2) software. Necrotic areas in H&E-stained sections and collagen deposition in Sirius red-stained slides were quantified from a minimum of three tumors and nine random field areas using the NIH ImageJ software (version 1.53e https://imagej.net/ij/download.html).

### Immunofluorescence

Slides were deparaffinized and rehydrated, and antigen retrieval was performed in R-BUFFER A (Electron Microscopy Sciences, 62706-10) in a Retriever (Electron Microscopy Sciences, 62700) for 1 h, followed by cooling at room temperature for 20 min. After blockade with 5% BSA in 0.1% Triton-X (Sigma, T8787), the tissue sections were incubated overnight with primary antibodies (Supplementary Table 1) at 4°C. The slides were washed and incubated with a fluorochrome-conjugated secondary antibody (Supplementary Table 2) for 1.5 h at room temperature in the dark. Following washing and nuclear staining with Hoechst (Invitrogen, H3570 1µg/mL in PBS) for 10 min, the slides were washed and mounted in DAKO fluorescent mounting medium (DAKO S3023) and dried for 3 h before visualization under a fluorescence microscope (Zeiss Axioscope 2). Quantification of positively stained area was performed using NIH Image J software. Immunofluorescence (IF) images were converted into red-green-blue (RGB) stacks. The threshold was corrected to count the cells in black, in contrast with the white background. The same threshold setting was set for all images.

### Flow cytometry

To profile the tumor infiltrating leukocytes (TILs), fresh tumor tissues collected at the endpoint were used. Tumor tissues were mechanically disrupted using a Tumor Dissociation Kit (Miltenyi Biotech, 130-096-730) according to the manufacturer’s protocol. Inguinal lymph nodes of the same mice, from the tumor draining side (DLN) and tumor non-draining contralateral side (NDLN), were also processed to obtain single-cell suspensions. TILs and DLN and NDLN cells were suspended in 2% FBS-PBS, counted and 1-2 ×10^5^ cells were stained with fluorochrome-conjugated antibodies (Supplementary Table 3). The cells were washed and analyzed by flow cytometry (CytoFLEX 30, Beckman Coulter). Data were analyzed using the FlowJo^TM^ software (BD Biosciences).

### Gene expression by RT-qPCR

Total RNA was extracted from the tumor tissues using the RNeasy Plus Mini Kit (Qiagen, #4134) following the manufacturer’s protocol. Two hundred micrograms of purified RNA was used to synthesize cDNA using the QuantiTect Reverse Transcription Kit (Qiagen, 205311). Quantitative RT-PCR was conducted for the indicated genes using the primers listed in Supplementary Table 4 and SsoAdvanced Universal SYBR Green Supermix (Bio-Rad, 1725274) in a CFX Connect real-time PCR detection system (Bio-Rad). Each primer was validated by melting and standard curve analyses. Cycle threshold (Ct) values were used to normalize gene expression levels between different samples and the housekeeping gene, *m36B4* (*Rplp0*). Fold induction was calculated by comparing the expression levels in B16-V tumors from *Nlrc5^+/+^* control mice.

### Immune cell depletion

To achieve CD8 T cell depletion, anti-CD8a monoclonal antibodies (clone 53-6.7, #BE0004-1, BioXCell) or an isotype control (Rat IgG2a, clone 2A3, #BE0089, BioXCell) were administered intraperitoneally at a dose of 25 μg per mouse on days −1 and +1 relative to tumor implantation. To deplete NK cells, anti-NK1.1 (PK136, #BE0036, BioXCell) or an isotype control was administered intraperitoneally at 200 μg per mouse per injection on days −1 and +1 prior to tumor implantation, with subsequent administrations every 7-8 days. Depletion of CD8 T cells and NK cells was confirmed 24 h post-administration by flow cytometry on lymph node and spleen cells in separate groups of mice.

### Gene expression analysis in TCGA dataset

Correlation between the expression of NLRC5 and candidate genes in the human Tumor Cancer Genome Atlas (TCGA) dataset on skin cutaneous melanoma (SKCM) was analyzed using the cBioPortal platform^74,75^.

### Statistics

Data analysis and graphic plotting were carried out using the GraphPad Prism (San Diego, CA, USA version 10.4.1). For other comparisons between groups, one-way ANOVA was used. *p* values were represented by asterisks: * <0.05, ** <0.01, *** <0.001, **** <0.001.

### Proteomic analysis

Protein samples were prepared from cell lines and snap frozen tumor tissues (20-50 mg), digested with trypsin, and the peptides collected as described in Supplementary methods. Liquid chromatography-tandem mass spectrometry (LC-MS/MS) analysis of peptides, protein identification and proteomic data analysis are detailed in Supplementary methods. Significantly modulated proteins with cut-off values of log2-fold change <−1 and > 1 and *p*-Value <0.05 in the FragPipeAnalystR pipeline output were sorted using Microsoft Excel (Office 365). GraphPad Prism version 10.0.3 (GraphPad, Boston, MA) was used to generate volcano plots and pie charts. Venn diagrams were generated using the jvenn online tool ^76^. The SRplot server ^77^ was used for Gene Ontology (GO) and pathway analyses and to generate enrichment plots and heatmaps.

## Data Availability

The mass spectrometry data are deposited to the ProteomeXchange Consortium via the PRIDE ^78^ partner repository with the dataset identifier PXD060523.

## Abbreviations

APC: antigen presenting cell
APM: antigen processing machinery
B16-N-FL: B16-F10 melanoma cells expressing NLRC5-FL
B16-N-SA: B16-F10 melanoma cells expressing NLRC5-SA
B16-V: B16-F10 melanoma cells expressing control vector
CC: cellular compartments
CTL: cytotoxic T lymphocytes
DC: dendritic cell
DEPs: differentially expressed proteins
DLN: draining lymph node
EC: endothelial cell
GO: gene ontology
ICB: immune checkpoint blockade
MF: molecular functions
MHC-I: MHC class-I
NDLN: non-draining lymph node
N-FL: NLRC5-FL full-length NLRC5
NLRC5-SA: NLRC5 super-activator
TILs: tumor infiltrating lymphocytes
VM: vascular mimicry.

## Funding and Acknowledgments

This work was funded by the Canadian Institutes of Health Research (CIHR) PJT-203990 to S.I and S.R. This research was enabled in part by support from the Centre de calcul scientifique de l’Université de Sherbrooke, Calcul Québec (calculquebec.ca) and the Digital Research Alliance of Canada (alliancecan.ca). CR-CHUS is an FRQS-funded research center. The authors thank Dr. Prenitha Doss for help in generating graphical abstract with Biorender.com. The authors thank Dr. Prenitha Doss for critical reading of the manuscript.

## Author Contributions

Conceptualization, S.I., S.R., T.A.K and A.S.

Methodology, A.S., A.J.I.Q., A.A.C., D.L. and S.R.;

formal analysis, A.S., J.-F.L., S.R. and S.I.

resources, F.-M.B., S.R. and S.I.

data curation, A.S., J.-F.L. and S.I.

writing-original draft preparation, A.S. and S.I.

writing-review and editing, A.S., D.L., J.-F.L., T.A.K., S.R. and S.I.

supervision, S.R. and S.I.

funding acquisition, S.R and S.I. All authors have read and agreed to the manuscript as a bioRxiv preprint.

## Ethics committee approval

All experiments on mice were carried out during daytime with the approval of the Université de Sherbrooke Ethics Committee for Animal Care and Use (Protocol ID 2023-4043).

