## Supplementary Information for "NLRC5 expression in tumor cells is critical to activate adaptive and innate antitumor immune responses"

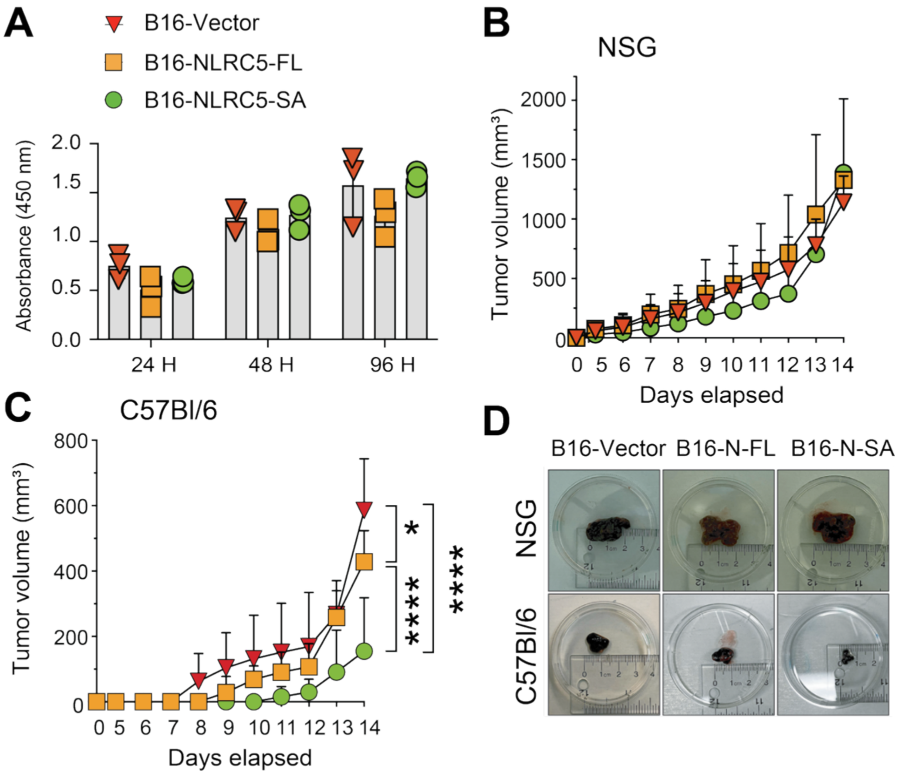


**Figure S1. Control of B16 melanoma expressing NLRC5 requires immune cells.**

(A) Expression of NLRC5 in tumor cells does not inhibit *in vitro* cell growth kinetics. B16-F10 melanoma cells expressing control vector (B16-V), full length NLRC5 (B16-N-FL) or engineered NLRC5 super-activator (B16-N-SA) were seeded in 96-well plates in triplicates and cell survival and growth was assessed by WST assay at the indicated time points.

(B, C) NLRC5 expressing B16 cells grow unhindered in immunocompromised *NOD.scid.gamma* (NSG) mice but inhibited in immune competent C57BL/6 mice. B16-V, B16-N-FL and B16-N-SA cell lines, $2 \times 10^5$ cells suspended in 50 µL PBS, were subcutaneously injected into the right flanks of 8 weeks-old NSG (B) or C57Bl/6 (C) mice. Tumor growth was monitored until B16-V tumors reached the endpoint in more than one of the NSG hosts. Calculated tumor volume from 4 mice per group.

(D) Representative images of tumors dissected from NSG and C57Bl/6 mice at the endpoint.

(A-C) Statistics: Mean + Standard deviation (SD). 2-way ANOVA with Tukey's multiple comparison test. * $p \leq 0.05$, **** $p \leq 0.0001$.

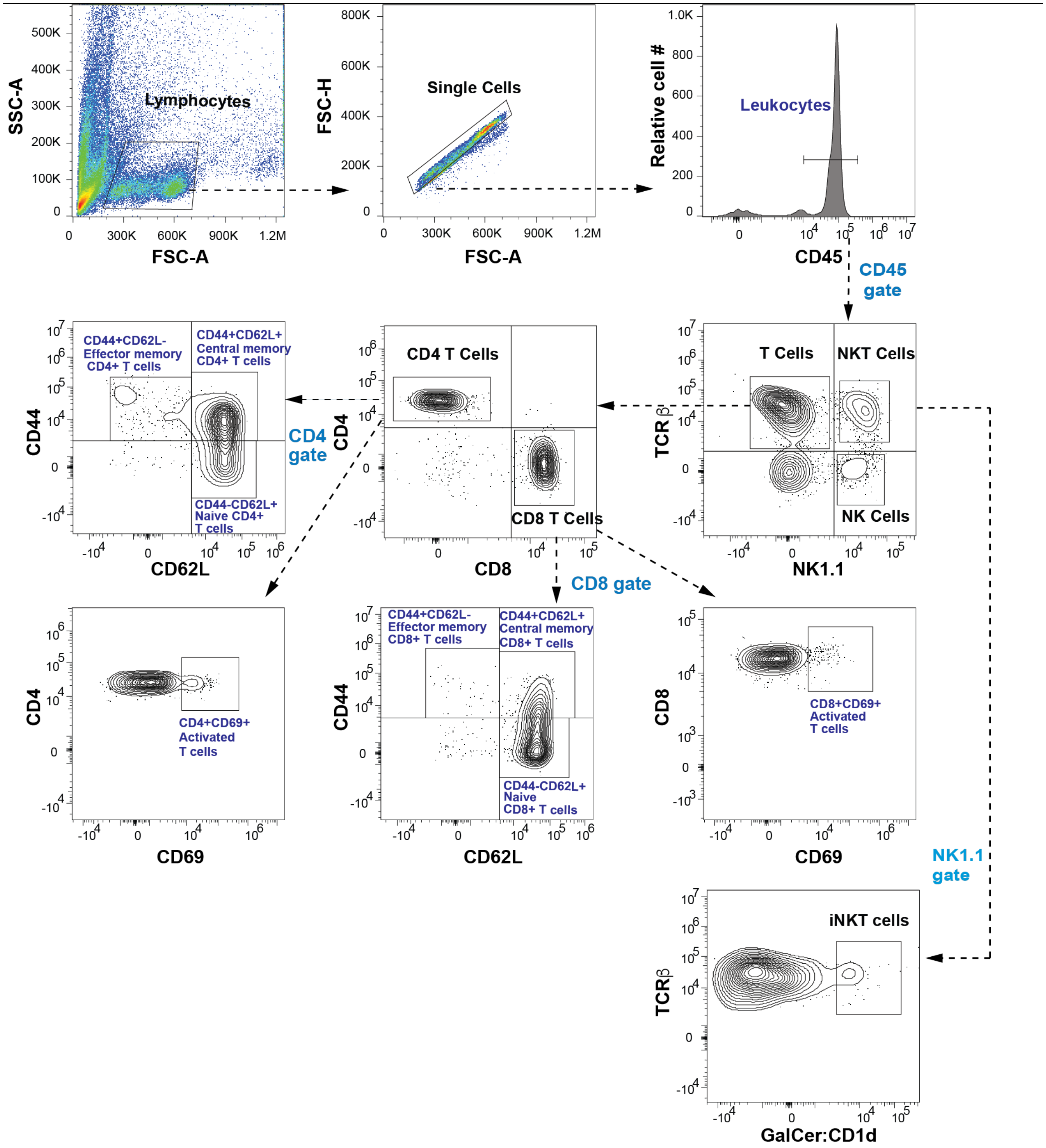


**Figure S2. Gating strategy to characterize T lymphocytes and NK cells infiltrating the NLRC5 expressing B16 tumors.** Tumor infiltrating lymphocytes (TILs) and single cell suspensions from tumor draining (DLN) and non-tumor draining (NDLN) lymph nodes were isolated at the time of tumor collection from *Nlrc5*$^{+/+}$ and *Nlrc5*$^{-/-}$ hosts implanted with B16-V, B16-N-FL or B16-N-SA tumors. Cells were counted and stained with fluorochrome conjugated antibodies (Supplementary Table 3) and analyzed by flow cytometry. Representative data from DLN are shown. $CD45^+$ leukocytes were gated from singlets, from which $CD3\varepsilon^+$ total T cells were identified and then grouped into $CD4^+$ and $CD8^+$ T lymphocytes. $CD4^+$ and $CD8^+$ T cells were sub-grouped into activated effector cells ($CD69^+$), effector memory cells ($CD44^+CD62L^-$), central memory ($CD44^+CD62L^+$) and naïve ($CD44^-CD62L^+$) cells. $CD45^+$ leukocytes expressing NK1.1 were gated and sub-grouped into NK, NKT and iNKT cells using TCRβ Ab and GalCer:CD1d reagent that binds the invariant Vα14Jα281 TCR in iNKT cells.

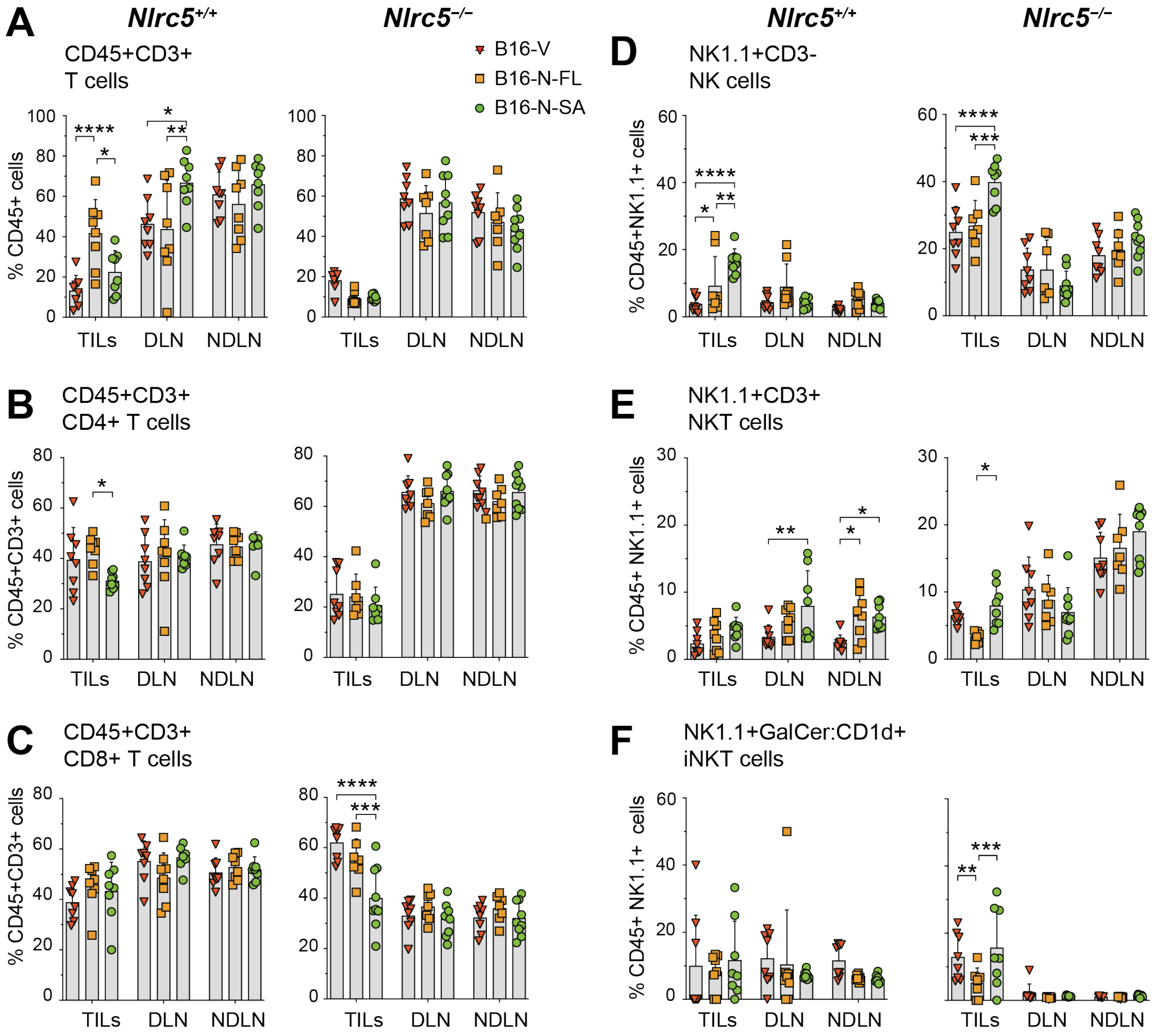
A
Nlrc5+/+
Nlrc5−/−
CD45+CD3+
T cells
B16-V
B16-N-FL
B16-N-SA
% CD45+ cells
TILs DLN NDLN
B
CD45+CD3+
CD4+ T cells
% CD45+CD3+ cells
C
CD45+CD3+
CD8+ T cells
% CD45+CD3+ cells
D
NK1.1+CD3-
NK cells
% CD45+NK1.1+ cells
E
NK1.1+CD3+
NKT cells
% CD45+ NK1.1+ cells
F
NK1.1+GalCer:CD1d+
iNKT cells
% CD45+ NK1.1+ cells

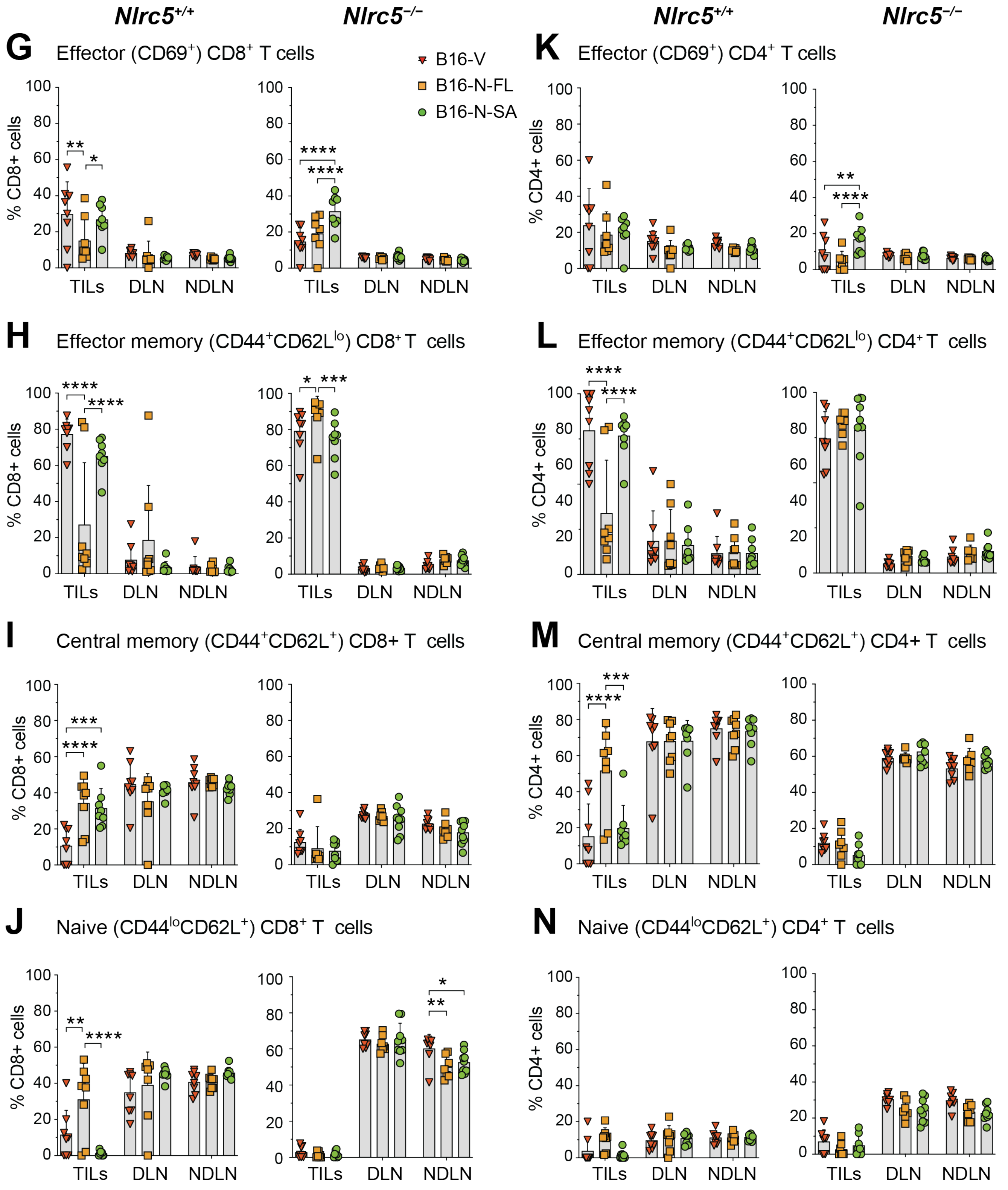


**Figure S3. Lymphoid and NK cell frequencies in NLRC5 expressing B16 tumors in *Nlrc5*$^{+/+}$ and *Nlrc5*$^{-/-}$ hosts.** TILs and single cell suspensions of DLNs and NDLNs were isolated from *Nlrc5*$^{+/+}$ and *Nlrc5*$^{-/-}$ hosts implanted with B16-V, B16-N-FL or B16-N-SA cells. Cells were counted and stained with fluorochrome conjugated antibodies for lymphoid and NK cell markers (Supplementary Table 3) and analyzed by flow cytometry using gating strategies depicted in Supplementary Fig. S2. Data from 6-10 mice per group from two independent experiments are shown.

(A-F) Proportions of $CD3^+$ T cells (A), $CD4^+$ T cells (B), $CD8^+$ T cells (C), NK cells (D, $NK1.1^+TCRb^-$), NKT cells (E, $NK1.1^+TCRb^+$) and iNKT cells (F, $NK1.1^+$ $TCRb^+$ αGalCer:$CD1d^+$).

(G-N) Proportions of $CD8^+$ and $CD4^+$ cells showing $CD69^+$ effector (G,K), $CD44^+CD62L^-$ effector memory (H, L), $CD44^+CD62L^+$ central memory (I, M) and $CD44^-CD62L^+$ naïve (J, N) phenotype.

Statistics: Mean + SD. Two-way ANOVA with Tukey's multiple comparison test. * $p \leq 0.05$, ** $p \leq 0.01$, *** $p \leq 0.001$, **** $p \leq 0.0001$. Significant values within B16-V, B16-N-FL and B16-N-SA are indicated by solid lines.

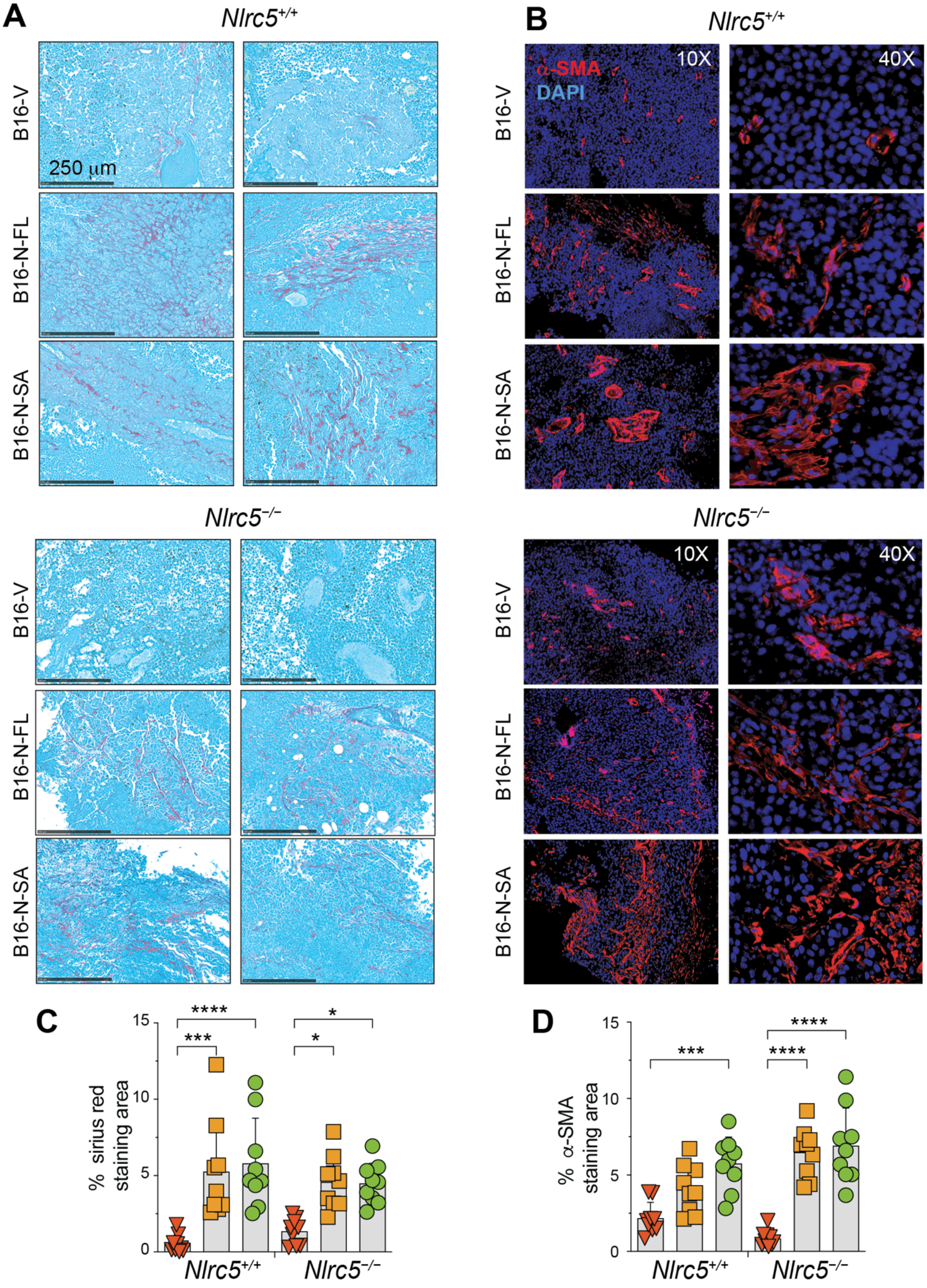


**Figure S4. B16-N-FL and B16-N-SA tumors display increased collagen deposition and myofibroblast content in *Nlrc5*$^{+/+}$ and *Nlrc5*$^{-/-}$ hosts**. (A,B) FFPE sections of B16-V, B16-N-FL and

B16-N-SA tumors resected from *Nlrc5*$^{+/+}$ and *Nlrc5*$^{-/-}$ hosts at the tumor endpoint were evaluated for (A) collagen deposition by Sirius red histochemical staining and (B) activated myofibroblasts by immunofluorescence (IF) staining of alpha smooth muscle actin (αSMA). Representative Sirius red staining areas from two different tumor sections for each group are shown (scale bar 250 µm). SMA staining is shown at 10× and 40× magnification. (C,D) The percentage of Sirius red and αSMA area was calculated from nine random fields from three tumors per group. Statistics: Mean + Standard deviation (SD). 2-way ANOVA with Tukey's multiple comparison test. * $p \leq 0.05$, *** $p \leq 0.001$, **** $p \leq 0.0001$.

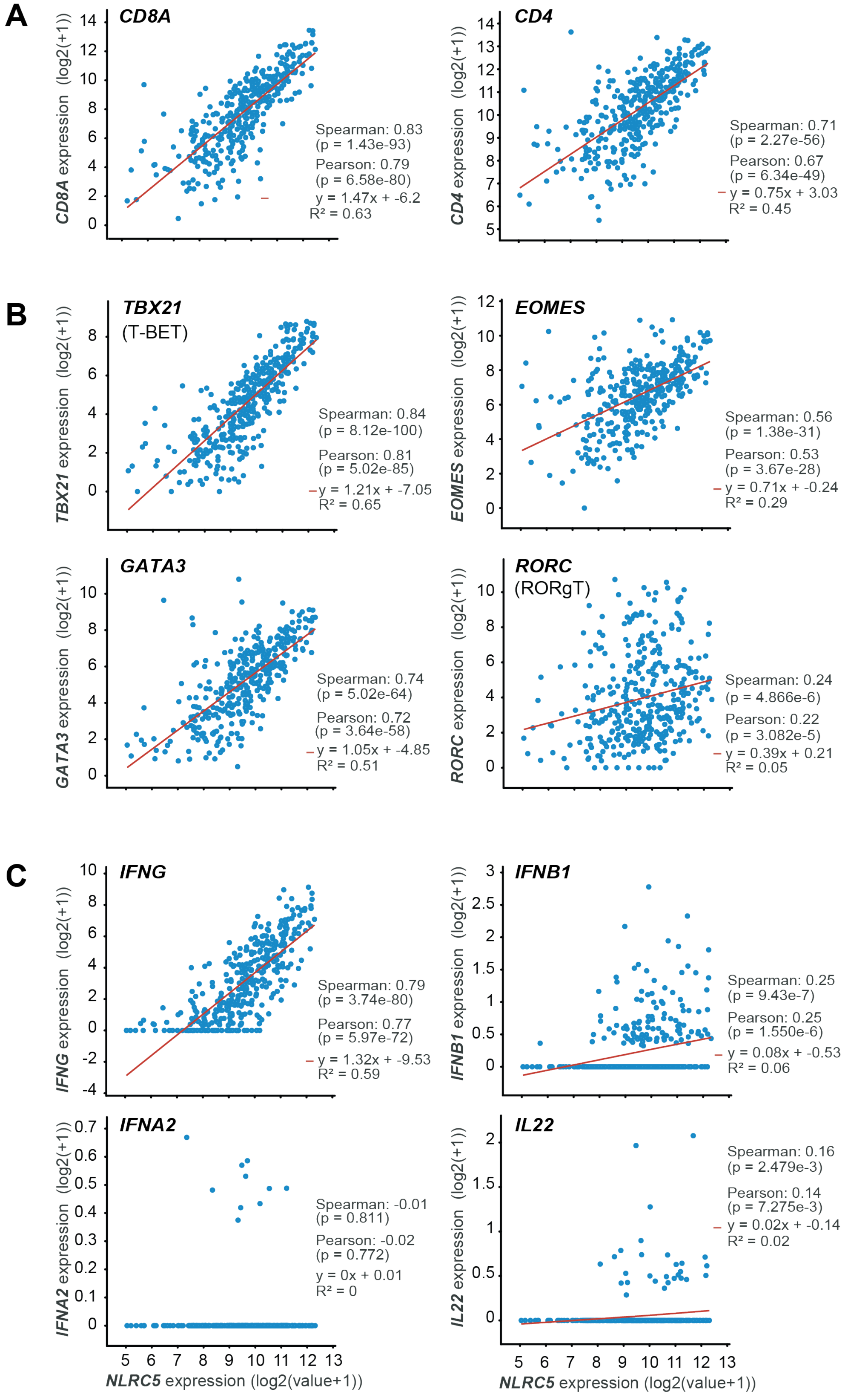


**Figure S5. NLRC5 expression in TCGA-SKCM positively correlates with T cell infiltration, expression of IFNγ and transcription factors regulating IFNγ.**

(A) Correlation between the expression of *NLRC5* and *CD8A* and *CD4* genes.

(B) Correlation between the expression of *NLRC5* and transcription factors regulating lymphoid cell differentiation and *IFNG* gene expression.

(C) Correlation between the expression of *NLRC5* and *IFN* genes.

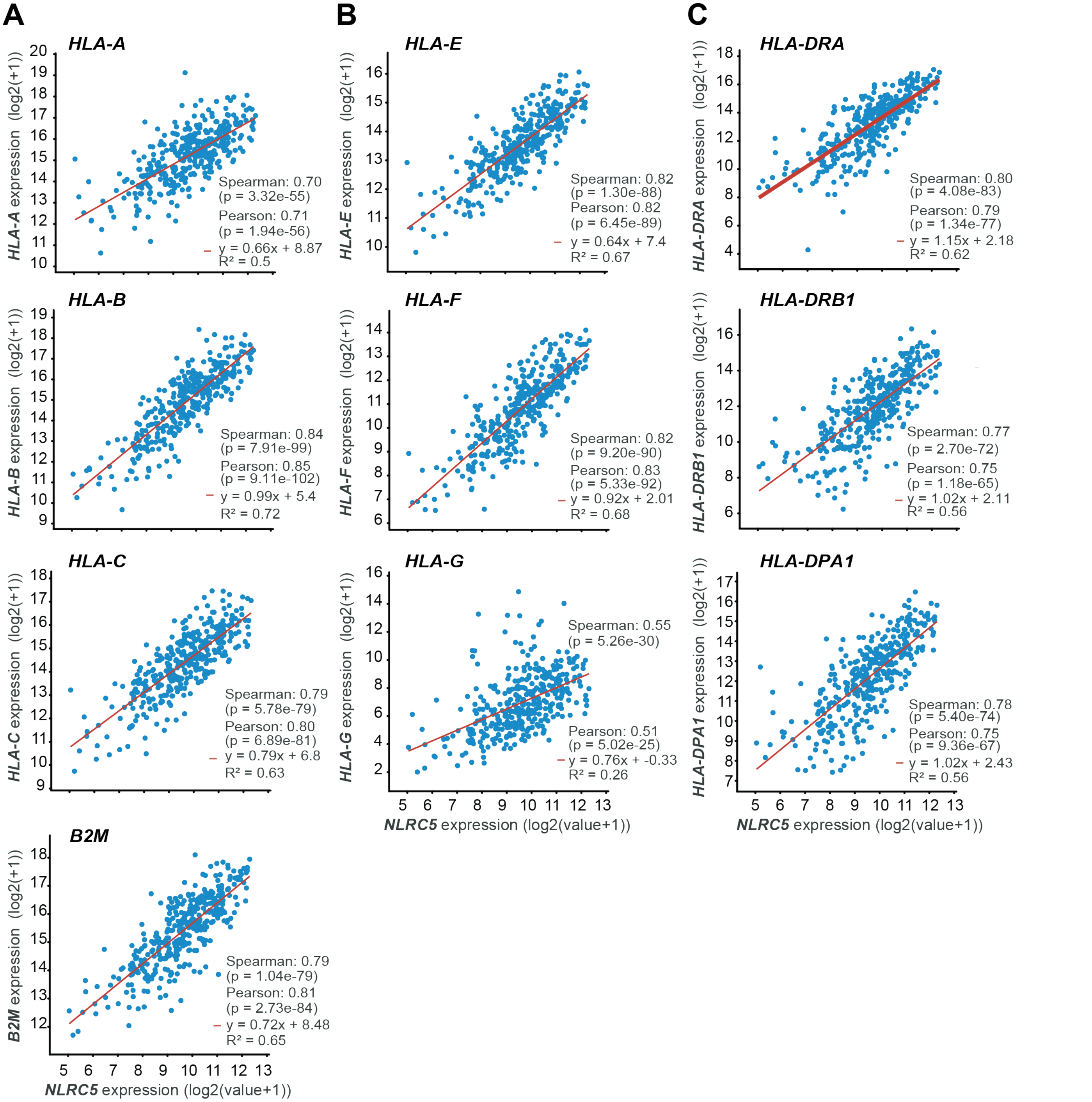


**Figure S6. NLRC5 expression in TCGA-SKCM positively correlates with MHC gene expression.**

(A) Correlation between the expression of *NLRC5* and classical MHC-Ia genes *HLA-A*, *HLA-B* and *HLA-C*, and *B2M*.

(B) Correlation between the expression of *NLRC5* and non-classical MHC-Ib genes *HLA-E*, *HLA-F* and *HLA-G*.

(C) Correlation between the expression of *NLRC5* and MHC-II genes *HLA-DRA*, *HLA-DRB1* and *HLA-DPA1*.

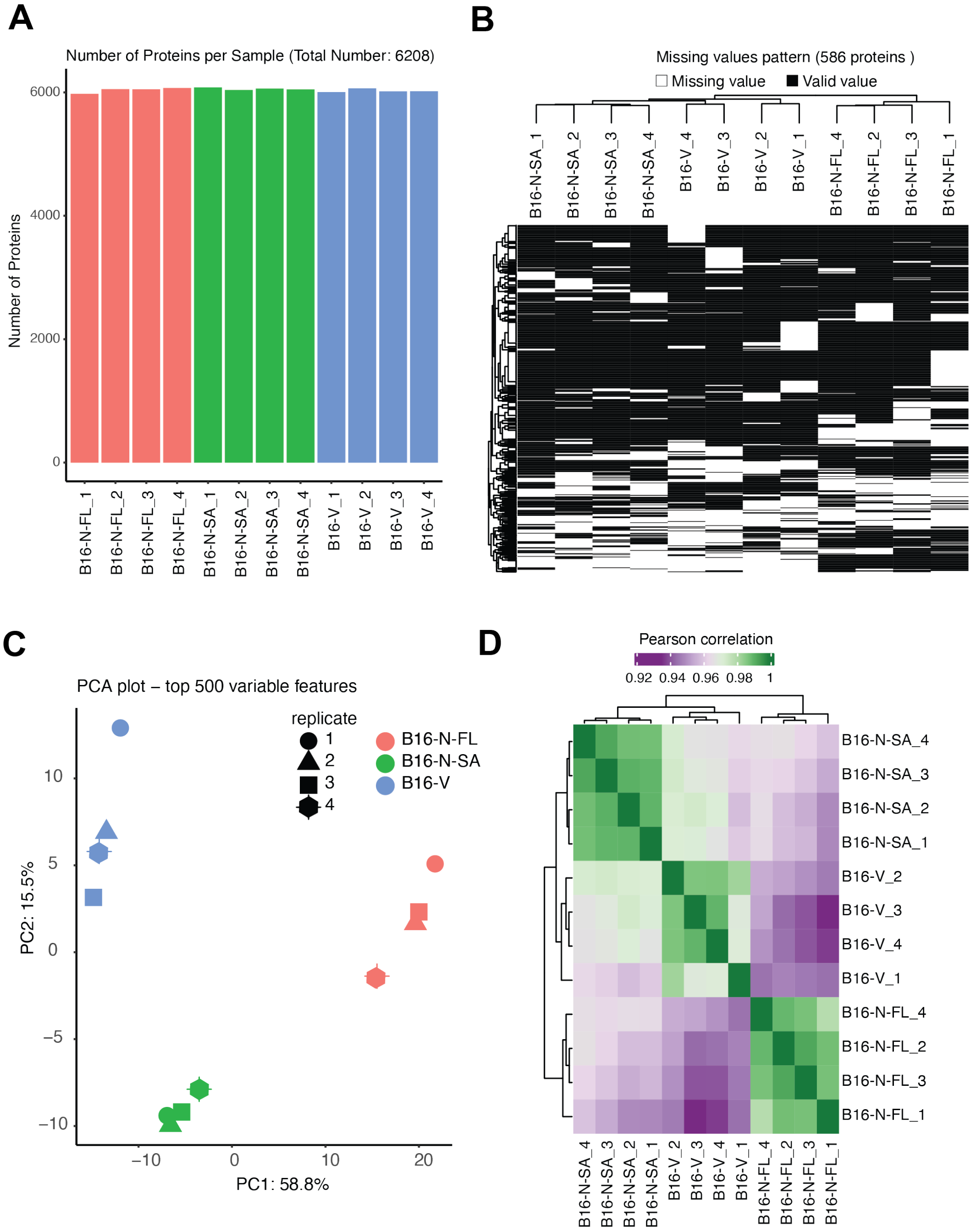


**Figure S7. Quality control for mass spectrometry data on tumor cell proteomes**. Quadruplicate samples of proteins extracted from B16-V, B16-N-FL and B16-N-SA cell lines were subjected to shotgun proteomics by LC-MS/MS. (A) Number of individual proteins identified in each tumor by more than one peptide. (B) Missing value pattern across samples. (C) Principal component analysis (PCA) plot. (D) Pearson correlation between samples.

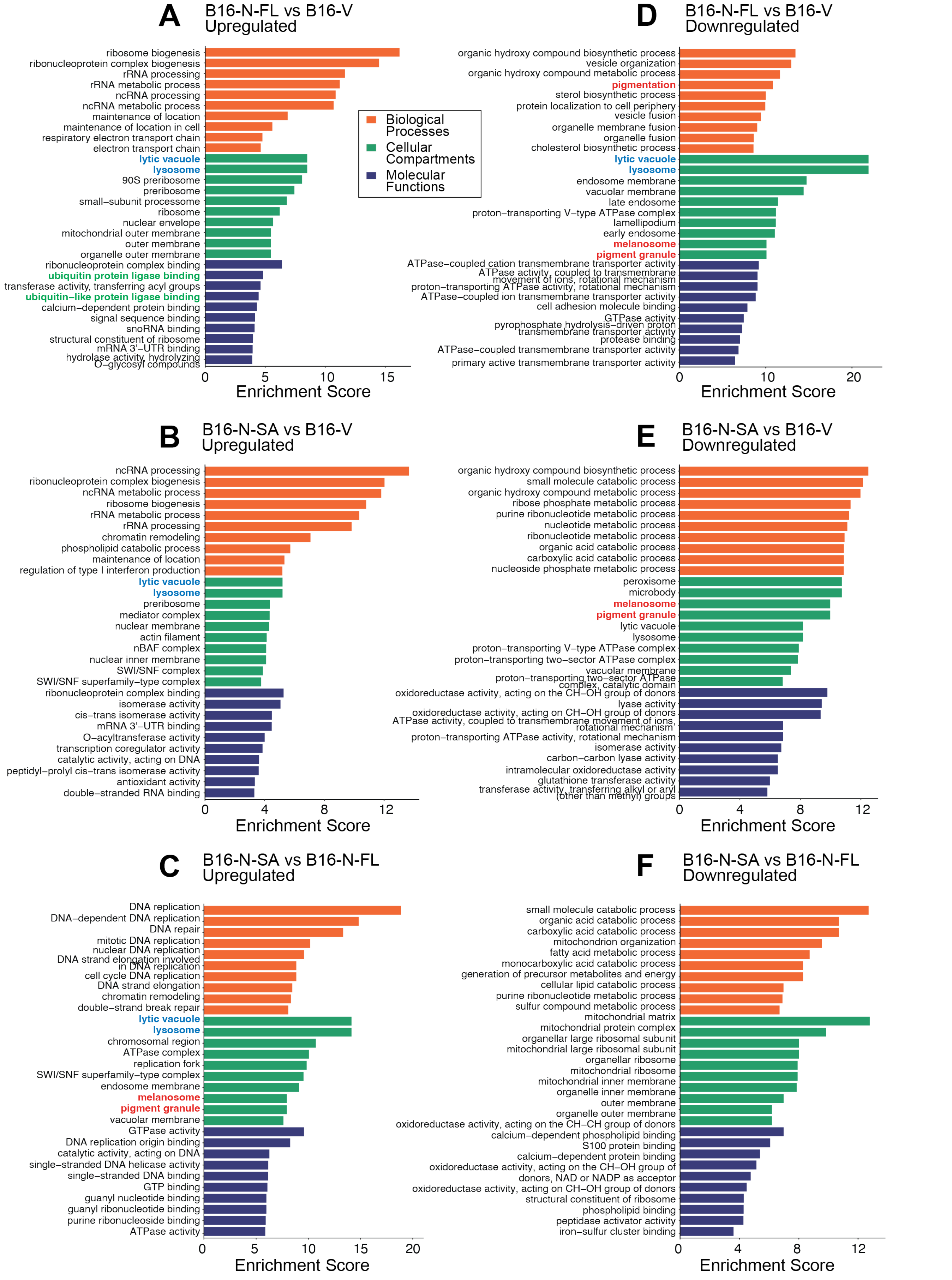
A
B16-N-FL vs B16-V
Upregulated
ribosome biogenesis
ribonucleoprotein complex biogenesis
rRNA processing
rRNA metabolic process
ncRNA processing
ncRNA metabolic process
maintenance of location
maintenance of location in cell
respiratory electron transport chain
electron transport chain
lytic vacuole
lysosome
90S preribosome
preribosome
small−subunit processome
ribosome
nuclear envelope
mitochondrial outer membrane
outer membrane
organelle outer membrane
ribonucleoprotein complex binding
ubiquitin protein ligase binding
transferase activity, transferring acyl groups
ubiquitin−like protein ligase binding
calcium−dependent protein binding
signal sequence binding
snoRNA binding
structural constituent of ribosome
mRNA 3'−UTR binding
hydrolase activity, hydrolyzing O−glycosyl compounds
0 5 10 15
Enrichment Score
Biological Processes
Cellular Compartments
Molecular Functions
D
B16-N-FL vs B16-V
Downregulated
organic hydroxy compound biosynthetic process
vesicle organization
organic hydroxy compound metabolic process
pigmentation
sterol biosynthetic process
protein localization to cell periphery
vesicle fusion
organelle membrane fusion
organelle fusion
cholesterol biosynthetic process
lytic vacuole
lysosome
endosome membrane
vacuolar membrane
late endosome
proton−transporting V−type ATPase complex
lamellipodium
early endosome
melanosome
pigment granule
ATPase−coupled cation transmembrane transporter activity
ATPase activity, coupled to transmembrane movement of ions, rotational mechanism
proton−transporting ATPase activity, rotational mechanism
ATPase−coupled ion transmembrane transporter activity
cell adhesion molecule binding
GTPase activity
pyrophosphate hydrolysis−driven proton transmembrane transporter activity
protease binding
ATPase−coupled transmembrane transporter activity
primary active transmembrane transporter activity
0 10 20
Enrichment Score
B
B16-N-SA vs B16-V
Upregulated
ncRNA processing
ribonucleoprotein complex biogenesis
ncRNA metabolic process
ribosome biogenesis
rRNA metabolic process
rRNA processing
chromatin remodeling
phospholipid catabolic process
maintenance of location
regulation of type I interferon production
lytic vacuole
lysosome
preribosome
mediator complex
nuclear membrane
actin filament
nBAF complex
nuclear inner membrane
SWI/SNF complex
SWI/SNF superfamily−type complex
ribonucleoprotein complex binding
isomerase activity
cis−trans isomerase activity
mRNA 3'−UTR binding
O−acyltransferase activity
transcription coregulator activity
catalytic activity, acting on DNA
peptidyl−prolyl cis−trans isomerase activity
antioxidant activity
double−stranded RNA binding
0 4 8 12
Enrichment Score
E
B16-N-SA vs B16-V
Downregulated
organic hydroxy compound biosynthetic process
small molecule catabolic process
organic hydroxy compound metabolic process
ribose phosphate metabolic process
purine ribonucleotide metabolic process
nucleotide metabolic process
ribonucleotide metabolic process
organic acid catabolic process
carboxylic acid catabolic process
nucleoside phosphate metabolic process
peroxisome
microbody
melanosome
pigment granule
lytic vacuole
lysosome
proton−transporting V−type ATPase complex
proton−transporting two−sector ATPase complex
vacuolar membrane
proton−transporting two−sector ATPase complex, catalytic domain
oxidoreductase activity, acting on the CH−OH group of donors
lyase activity
oxidoreductase activity, acting on CH−OH group of donors
ATPase activity, coupled to transmembrane movement of ions, rotational mechanism
proton−transporting ATPase activity, rotational mechanism
isomerase activity
carbon−carbon lyase activity
intramolecular oxidoreductase activity
glutathione transferase activity
transferase activity, transferring alkyl or aryl (other than methyl) groups
0 4 8 12
Enrichment Score
C
B16-N-SA vs B16-N-FL
Upregulated
DNA replication
DNA−dependent DNA replication
DNA repair
mitotic DNA replication
nuclear DNA replication
DNA strand elongation involved in DNA replication
cell cycle DNA replication
DNA strand elongation
chromatin remodeling
double−strand break repair
lytic vacuole
lysosome
chromosomal region
ATPase complex
replication fork
SWI/SNF superfamily−type complex
endosome membrane
melanosome
pigment granule
vacuolar membrane
GTPase activity
DNA replication origin binding
catalytic activity, acting on DNA
single−stranded DNA helicase activity
single−stranded DNA binding
GTP binding
guanyl nucleotide binding
guanyl ribonucleotide binding
purine ribonucleoside binding
ATPase activity
0 5 10 15 20
Enrichment Score
F
B16-N-SA vs B16-N-FL
Downregulated
small molecule catabolic process
organic acid catabolic process
carboxylic acid catabolic process
mitochondrion organization
fatty acid metabolic process
monocarboxylic acid catabolic process
generation of precursor metabolites and energy
cellular lipid catabolic process
purine ribonucleotide metabolic process
sulfur compound metabolic process
mitochondrial matrix
mitochondrial protein complex
organellar large ribosomal subunit
mitochondrial large ribosomal subunit
organellar ribosome
mitochondrial ribosome
mitochondrial inner membrane
organelle inner membrane
outer membrane
organelle outer membrane
oxidoreductase activity, acting on the CH−CH group of donors
calcium−dependent phospholipid binding
S100 protein binding
calcium−dependent protein binding
oxidoreductase activity, acting on the CH−OH group of donors, NAD or NADP as acceptor
oxidoreductase activity, acting on CH−OH group of donors
structural constituent of ribosome
phospholipid binding
peptidase activator activity
iron−sulfur cluster binding
0 4 8 12
Enrichment Score

**Figure S8. Gene ontology analysis of differentially expressed proteins in B16 tumor cell lines**. Gene ontology (GO) analysis of upregulated (A-C) and downregulated (D-F) proteins in B16-N-FL cells compared to B16-V cells (A,D), B16-N-SA versus B16-V cells (B,E), and B16-N-SA versus B16-N-FL cells (C,F); vs, versus. GO terms in Biological Processes (BP, orange bars), Cellular Compartments (CC, green bars), and Molecular Functions (MF, blue bars) are indicated with corresponding enrichment scores. GO terms that are upregulated in B16-N-FL and B16-N-SA cells (CC: lysosome and lytic vacuole), downregulated in B16-N-FL and B16-N-SA cells (CC: melanosome, pigment granule), upregulated in B16-N-FL cells (MF: ubiquitin protein ligase, ubiquitin-like protein ligase) are highlighted by colored fonts.

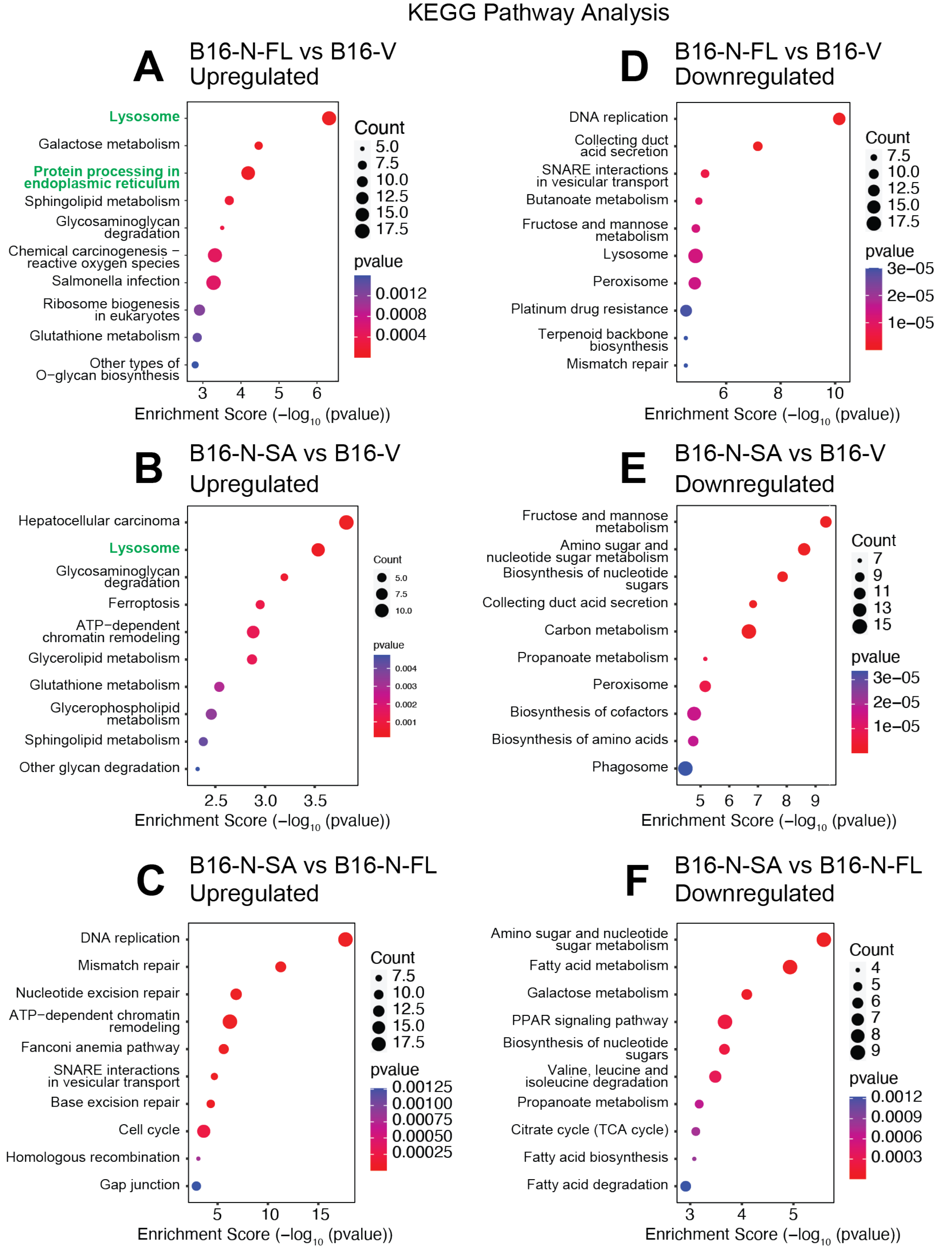


**Figure S9. Pathway analysis of differentially expressed proteins in B16 tumor cell lines**. Pathway analysis of upregulated (A-C) and downregulated (D-F) proteins in B16-N-FL cells compared to B16-V cells (A,D), B16-N-SA versus B16-V cells (B,E), and B16-N-SA versus B16-N-FL cells (C,F). Pathway terms upregulated in B16-N-FL and B16-N-SA cells (lysosome), and upregulated in B16-N-FL (protein processing in the endoplasmic reticulum) are highlighted by colored fonts.

***Proteomic analysis***

***Protein preparation and protease digestion***

Snap frozen tumor tissues (20-50 mg) were resuspended in 1 mL of lysis buffer (8 M Urea, 1 M $NH_4HCO_3$ and 10 mM HEPES-KOH pH7.5) in a 2 mL low protein binding tubes (Axygen). Tissues were homogenized using steel beads in a mixer mill (MM 400, Retsch, Haan, Germany). Tissue lysates were transferred to fresh tubes and sonicated on ice (12 cycles, 20-25% intensity, 5 sec PULSE/ 5 sec OFF) and centrifuged at 16,000 *g* for 10 min at 4°C. Supernatants were transferred to fresh tubes and proteins quantified using DC Protein Assay Kit (Bio-Rad, #5000113, #5000114, #5000115) following the manufacturer's instructions. One hundred µg of protein was transferred to new tubes, volumes adjusted to 50 µL with urea solution (8 M Urea, 10 mM HEPES, pH 8) and 1 µL of 255 mM dithiothreitol was added to achieve 5 mM final concentration. The tubes were vortexed and boiled at 95°C for 2 min and allowed to cool at room temperature for 30 min. After adding 1.5 µL of photosensitive 262.5 mM chloroacetamide to achieve 7.5 mM final concentration, the samples were vortexed and incubated in dark at room temperature for 20 min before adding 150 µL of 50 mM $NH_4HCO_3$ to bring the urea concentration down to 2 M. Proteins were digested by adding 1 µg of Pierce™ Trypsin Protease (Thermo Scientific, cat# 90058) and overnight incubation at 30°C. Proteolysis was stopped by adding 0.5 µL of 100% Trifluoroacetic acid (TFA), and the digested peptides were cleaned using Pierce™ C18 tips (Thermo Scientific, cat# 87784). The solvents were removed using a Vacufuge Plus centrifuge concentrator (Eppendorf), and the peptides diluted in 1% Formic Acid (FA) were quantified using a NanoDrop 2000/2000c spectrophotometer (Thermo Fisher Scientific).

***Liquid chromatography-tandem mass spectrometry (LC-MS/MS)***

Concentrated peptides (250 ng) were separated using a nanoHPLC system (nanoElute, Bruker Daltonics). The samples were loaded onto an Acclaim PepMap100 C18 Trap Column (0.3 mm id x 5 mm, Dionex Corporation) at a consistent flow of 4 µL/min, and peptides were eluted onto a PepMap C18 analytical nanocolumn (1.9 µm bead size, 75 µm x 25 cm, PepSep) heated at 50°C. Peptides were eluted with solvent B (100% ACN and 0.1% FA) in a 5-37% linear gradient at a flow rate of 400 nL/min for ~2 h. The HPLC system was coupled to an TimsTOF Pro ion mobility mass spectrometer containing a Captive Spray nanoelectrospray source (Bruker Daltonics) was used. Data acquisition was performed using the diaPASEF mode. For each individual Trapped Ion Mobility Spectrometry (TIMS) measurement in diaPASEF mode, a single mobility window consisting of 27 mass steps (with m/z ranging from 114 to 1414 and a mass width of 50 Da) was employed per cycle, which had a 1.27-second duty cycle. This process involves scanning the diagonal line in the m/z-ion mobility plane for +2 and +3 charged peptides.

***Protein identification***

Peptide mass spectra were analyzed using the DIA-NN , an open-source software [1] suite for DIA / SWATH data processing (https://github.com/vdemichev/DiaNN, version 1.8.1), installed in an Apptainer container (https://apptainer.org/, version 1.3.5) using docker image provided on the docker hub [2]. Analysis was performed using default parameters, except for the following options: two missed cleavages were allowed; trypsin digestion was performed for K/R; and protein N-term methionine excision was used as a variable modification for the *in-silico* digest.

The *Mus musculus* reference proteome UP000000589 was downloaded from the UniProt website (https://www.uniprot.org/proteomes/UP000000589). The reference proteome contained 63367 proteins. For the FASTA search, DIA-NN was instructed to perform an *in silico* digest of the sequence database. A mass tolerance accuracy of MS1 and MS2 of 20 ppm was used for the precursor and fragment ions, respectively. Minimum and maximum values were set for peptide length (7-30 amino acids), precursor charge (1-5), precursor m/z (100-1700) and fragmentation m/z (100-1500) for *in silico* library generation or library-free search. For re-analysis, match between runs (MBR) was enabled and smart profiling was chosen when creating a spectral library from DIA data. Carboxyamidomethylation (unimod4) and oxidation (M) (unimod35) were set as fixed modifications, and N-terminal protein acetylation was set as a variable modification. DIANN protein group matrix was filtered using a custom Perl script to extract protein groups with a single protein. This new matrix was used as input to the R package FragPipeAnalystR [3] to obtain QC metrics and differential protein expression profiles. Significantly modulated proteins with cut-off values of log2-fold change <-1 and >1 and *p*-Value <0.05 in the FragPipeAnalystR pipeline output were sorted using Microsoft Excel (Office 365). GraphPad Prism was used to generate volcano plots and pie charts. Venn diagrams were generated using the jvenn online tool [4]. The SRplot server [5] was used for Gene Ontology (GO) and pathway analyses and to generate enrichment plots and heatmaps.

**Supplementary Table 1: Primary antibodies and reagents used for IF**

| Antibody | Species | Source | Catalogue # | Dilution |
|---|---|---|---|---|
| CD45 | Rat | Cell Signaling | 30-F11 | 1:250 |
| CD31 | Goat | R&D System | AF3628 | 1:250 |
| αSMA | Mouse | Abcam | ab7817 | 1:150 |
| Tomato Lectin (TL) | Rodents | VectorLabs | DL-1174-1 | 1:250 |

**Supplementary Table 2: Secondary antibodies used for IF**

| Antibody | Source | Catalogue # | Dilution |
|---|---|---|---|
| Alexa Fluor 488 goat Anti-rat IgG (H&L) | Invitrogen | A-11006 | 1:500 |
| Alexa Fluor 568 goat Anti-mouse IgG (H&L) | Invitrogen | A-11031 | 1:500 |
| Alexa Fluor 488 horse Anti-goat IgG (H&L) | VectorLabss | DI-3088 | 1:500 |
| Alexa Fluor 594 horse Anti-goat IgG (H&L) | VectorLabs | DI-3094 | 1:500 |

**Supplementary Table 3: List of antibodies used for flow cytometry**

| Marker | Fluorochrome | Clone | Source | #Cat |
|---|---|---|---|---|
| CD45 | Brilliant Violet 605 | 30-F11 | Biolegend | 103140 |
| CD3e | Brilliant Violet 510 | 145-2C11 | Biolegend | 100353 |
| TCR β chain | PE/Dazzle 594 | 109240 | Biolegend | H57-597 |
| CD4 | Alexa Flour 700 | GK1.5 | ebiosciences | 5016851 |
| CD8a | eFLOUR 450 | 53-6.7 | ebiosciences | 48-0081-82 |
| CD62L | APC | Mel-14 | ebiosciences | 17-0621-83 |
| CD44 | FITC | IM7 | Biolegend | 103006 |
| CD69 | PRCy7 | H1.2F3 | Biolegend | 104512 |
| B220/CD45R | PerCP | RA3-6B2 | Biolegend | 103233 |
| NK1.1 | APC-Cy7 | PK136 | Biolegend | 108724 |
| α-GalCer:CD1d | PE | L363 | Biolegend | 140506 |

**Supplementary Table 4: RT-qPCR primer sequences**

| Gene | Accession # | Forward Primer | Reverse Primer | Amplic-on (bp) |
|---|---|---|---|---|
| *Klrb1c* | NM_008527 | TGGCACAACTTTCAATTCTGATG | GGTTTAGTTCCTTTTGGCAGATC | 140 |
| *Ncr1* | NM_010746 | TCTGGTCAAAGTCGAGCAAC | TGTGGCAGTCTTCAGTTGG | 137 |
| *Itga2* | NM_008396 | CGATACACATAACCCTCAGCTC | CTGCCTATGATAACCCCTGTC | 114 |
| *Klrk1* | NM_001083322 | GCTGGTTAAGTCCTATCACTGG | TTGAGCCATAGACAGCACAG | 143 |
| *Klrc2* | NM_010653 | AATCTCTTTCACTGGTCTCATGG | GCAAATTCATCTAAAGGGAGCC | 119 |
| *Klrd1* | NM_010654 | TCAACACCTTCTCCAACCAC | CTGATGCCCAACCCACTT | 84 |
| *Klra10* | NM_008459 | AGAACAGGACAGATGGAATAGTG | TCGCTTTACATCCACTCCATG | 146 |
| *Rae1* | NM_175112 | CTCTGGATTTGGGACTGGTG | GGAAGTTGCCTGGTAAAGTTG | 149 |
| *Ccl2* | NM_011333 | GTCCCTGTCATGCTTCTGG | GCTCTCCAGCCTACTCATTG | 144 |
| *Ccl4* | NM_013652 | AAACCTAACCCCGAGCAAC | CGGGAGGTGTAAGAGAAACAG | 144 |
| *Ccl5* | NM_013653 | GGGTACCATGAAGATCTCTGC | TCTAGGGAGAGGTAGGCAAAG | 129 |
| *Ccl17* | NM_011332 | AGACCTTCACCTCAGCTTTTG | CTTTGAAGTAATCCAGGCAGC | 144 |
| *Cxcl2* | NM_009140 | AATGCCTGAAGACCCTGC | TTTTGACCGCCCTTGAGAG | 118 |
| *Cxcl5* | NM_009141 | GTTCCATCTCGCCATTCATG | TTAAGCAAACACAACGCAGC | 129 |
| *Cxcl9* | NM_008599 | AGTCCGCTGTTCTTTTCCTC | TGAGGTCTTTGAGGGATTTGTAG | 141 |
| *Cxcl10* | NM_021274 | TCAGCACCATGAACCCAAG | CTATGGCCCTCATTCTCACTG | 144 |
| *Cxcl11* | NM_019494 | ATGGCAGAGATCGAGAAAGC | TGCATTATGAGGCGAGCTTG | 137 |
| *Cxcl12* | NM_001012477 | ACTCCAAACTGTGCCCTTC | AAGCTTTCTCCAGGTACTCTTG | 106 |
| *Ifna* | NM_010502 | TCTGTGCTTTCCTGATGGTC | GGTTATGAGTCTGAGGAAGGTC | 84 |
| *Ifnb* | NM_010510 | CAGCCCTCTCCATCAACTATAAG | TCTCCGTCATCTCCATAGGG | 144 |
| *Ifng* | NM_008337 | CCTAGCTCTGAGACAATGAACG | TTCCACATCTATGCCACTTGAG | 150 |
